# Tau filaments are tethered within brain extracellular vesicles in Alzheimer’s disease

**DOI:** 10.1101/2023.04.30.537820

**Authors:** S. L. Fowler, T. S. Behr, E. Turkes, P. Maglio Cauhy, M. S. Foiani, A. Schaler, G. Crowley, S. Bez, E. Ficulle, E. Tsefou, D. P. O’Brien, R. Fischer, B. Geary, P. Gaur, C. Miller, P. D’Acunzo, E. Levy, K. E. Duff, B. Ryskeldi-Falcon

## Abstract

The abnormal assembly of tau protein in neurons is the pathological hallmark of multiple neurodegenerative diseases, including Alzheimer’s disease (AD). In addition, assembled tau associates with extracellular vesicles (EVs) in the central nervous system of patients with AD, which is linked to its clearance and prion-like propagation between neurons. However, the identities of the assembled tau species and the EVs, as well as how they associate, are not known. Here, we combined quantitative mass spectrometry, cryo-electron tomography and single-particle cryo-electron microscopy to study brain EVs from AD patients. We found filaments of truncated tau enclosed within EVs enriched in endo-lysosomal proteins. We observed multiple filament interactions, including with molecules that tethered filaments to the EV limiting membrane, suggesting selective packaging. Our findings will guide studies into the molecular mechanisms of EV-mediated secretion of assembled tau and inform the targeting of EV-associated tau as potential therapeutic and biomarker strategies for AD.

## INTRODUCTION

Alzheimer’s disease (AD) is characterised by intraneuronal inclusions of phosphorylated assembled tau protein, called neurofibrillary tangles, and extracellular plaques of assembled amyloid-β peptides. Tau assembles into two types of amyloid filament in neurofibrillary tangles, paired helical filaments (PHFs) and straight filaments (SFs), which are formed of two identical protofilaments with a C-shaped fold^1, 2^. Tau assembly correlates with neuronal dysfunction and clinical symptoms in AD^3, 4^, as well as in a large group of neurodegenerative diseases collectively known as tauopathies. Rare mutations in the tau gene, *MAPT*, lead to tau assembly and frontotemporal dementia^5–7^, demonstrating a causal role of tau assembly in neurodegeneration.

In AD, tau assembly begins in the locus coeruleus and transentorhinal cortex, and progresses over years to synaptically-connected regions in the limbic system and neocortex^8, 9^. Studies of tau assembly in cell and *in vivo* models suggest that prion-like propagation may underlie this progression, including the intracellular trafficking, intercellular transfer and seeded aggregation of assembled tau^10–13^.

Extracellular vesicles (EVs) are lipid membrane-enclosed compartments that are released and taken up by many cell types, including neurons and glia^14^. They have been shown to selectively shuttle proteins, RNA, lipids and metabolites between cells^15^, and have been implicated in synaptic transmission in neurons^16, 17^. EVs can be broadly categorised as exosomes or microvesicles (also known as ectosomes), based on their biogenesis pathway^15^. Exosomes are generated by budding of the endosomal membrane into the lumen of secretory multivesicular endosomes (MVEs) and are released into the extracellular space upon fusion of MVEs with the plasma membrane. Microvesicles bud directly from the plasma membrane. EVs can enter into recipient cells by endocytosis and deliver their contents to the cytoplasm by fusion with the limiting membranes of endo-lysosomes^15, 18, 19^.

Phosphorylated insoluble tau is associated with EVs isolated from AD patient brain and cerebrospinal fluid^20–23^. Similar observations have been made for EVs from transgenic mice expressing mutant tau^24^. EVs from the central nervous system of AD patients seed tau assembly when added to cultured cells and when injected into the brains of tau transgenic mice^21–23, 25^. EVs deliver assembled tau to the cytoplasm of recipient cells via the endo- lysosomal system^26^. Studies comparing non-EV-associated and EV-associated assembled tau have shown that EV-associated tau is more potent at seeding assembly^22, 23, 27, 28^. These studies suggest that EVs have the potential to mediate the prion-like propagation of assembled tau in human brain.

The molecular species and structures of assembled tau, and the identities of the EVs that they associate with are not known. In addition, how assembled tau associates with EVs, including if it is luminal or surface-bound, is unclear. This information would provide insights into the molecular mechanisms governing the secretion of assembled tau within EVs. It will also inform AD biomarker discovery and therapeutic strategies targeting assembled tau.

Here, we isolated EVs from AD patient brain using density gradient centrifugation and profiled their protein composition using quantitative mass spectrometry. We confirmed the ability of the EVs to seed tau assembly in cultured cells and *in vivo*. We characterised the molecular species and structures of assembled tau associated with EVs using quantitative mass spectrometry, biochemical fractionation and cryo-EM. Finally, we used cryo-ET to investigate the association of assembled tau with EVs. Our results show that PHFs and SFs formed of truncated tau are tethered inside EVs enriched in endo-lysosomal proteins in AD patient brain.

## RESULTS

### Isolation of EV populations from AD patient brain

We isolated EVs from flash-frozen post-mortem AD patient brain tissue across eight fractions using density gradient centrifugation (Figure 1 and Table S1). Immunoblotting and quantitative mass spectrometry of EV fractions isolated from frontal cortex, temporal cortex and hippocampus demonstrated the enrichment of known EV-associated proteins, including the microvesicle-enriched proteins annexin A2 and flotillin 1, the exosome-enriched tetraspanin CD81 and the lysosomal-associated membrane protein 2 (LAMP2) (Figure 1A)^29^. Intracellular proteins, such as lamin A/C, were depleted. We confirmed the ability of the EVs to seed tau assembly in the Tau RD P301S FRET Biosensor cell line (Figures S1A and S1B)^30^, as well in transgenic mice expressing human P301S tau (PS19 line)^31^ two months following injection into the hippocampal dentate gyrus (Figures S1C and S1D).

**Figure 1:**
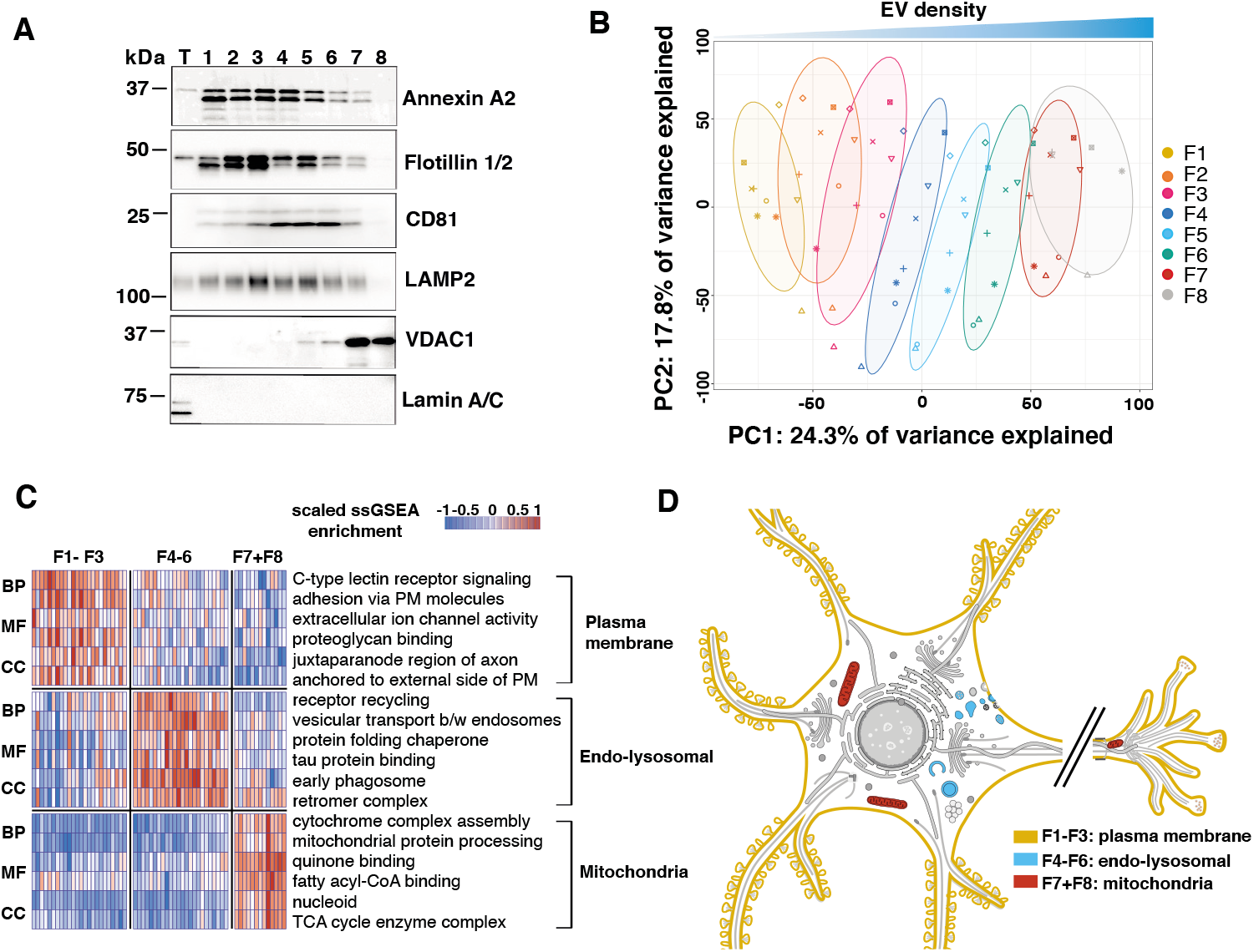
Proteomic characterisation of extracellular vesicles from Alzheimer’s disease patient brain. (A) Representative immunoblot of total homogenate (T) and EV fractions 1–8 isolated from AD patient brain using antibodies against annexin A2, flotillin 1, CD81, LAMP2, VDAC1 and lamin A/C. n=5. (B) Principal component (PC) analysis of quantitative mass spectrometry of EV fractions 1–8 from human AD brain. Symbols denote different cases. n=8 for F1–7 and n=5 for F8. Ellipses indicate 68% confidence intervals. (C) Gene ontology (GO) pathway enrichment analysis of quantitative mass spectrometry of grouped EV fractions 1–3, 4–6 and 7–8. Each column represents an individual case. n=8. Per row scaled ssGSEA enrichment scores are shown. BP: biological process, MF: molecular function, CC: cellular component. (D) Enriched biosynthetic subcellular localisation of enriched proteins in each EV group using the SwissBioPics animal neuronal cell.

Nanoparticle tracking analysis (NTA) showed that the majority of EVs were collected in the lightest density fraction 1 (Figure S2A). EVs in fractions 1–7 had similar size distributions, with average mode diameters of 132.1 nm ± 13.98 nm. In contrast, EVs in fraction 8 were significantly larger, with diameters of 163.5 nm ± 19.17 nm (Figures S2B–S2D). Across all fractions, 10% of EVs had diameters smaller than 92.5 nm (D10) and 10% had diameters larger than 262.5 nm (D90) (Figure S2C).

We developed a novel imputation strategy for the detection of enriched proteins amongst fractions in our quantitative mass spectrometry analysis, and detected a total of 6,105 proteins. Gene ontology (GO) and pathway enrichment analyses revealed marked differences in the protein compositions of the EV fractions (Figures 1B–1D). Fractions 1–3 were enriched in plasma membrane-associated proteins, suggesting a predominance of microvesicles. In contrast, fractions 4–6 were enriched in endo-lysosomal proteins, suggesting they mainly contained exosomes. Fractions 7 and 8 were enriched in mitochondrial proteins, supported by immunoblotting showing that the mitochondrial voltage-dependent anion channel 1 was restricted to these fractions (Figure 1A). EVs in fractions 7 and 8 are consistent with recently-described mitovesicles^32^. These results demonstrate that EV populations enriched in distinct organelle components can be isolated from post-mortem AD patient brain tissue based on density.

### EVs from AD patient brain are associated with assembled truncated tau

We investigated the association of tau with the EV populations from AD patient brain in our quantitative mass spectrometry dataset. Peptides encompassing the third and fourth tau repeats (residues 322-370) were the most enriched. Peptides from the extreme N-terminus (residues 6-23), the other two repeats (residues 242–254 and 281–311) and the C-terminal half of the proline-rich region (residues 212–224) were also enriched (Figures 2A and 2B). In agreement with these results, immunoblotting of the EV fractions showed that the majority of tau existed as approximately 12 and 24 kDa truncated species that were detected by an antibody against the third and fourth tau repeats (TauC), but not by antibodies against epitopes between the extreme N-terminus and the proline-rich region (Tau13 and HT7) (Figure 2C).

**Figure 2:**
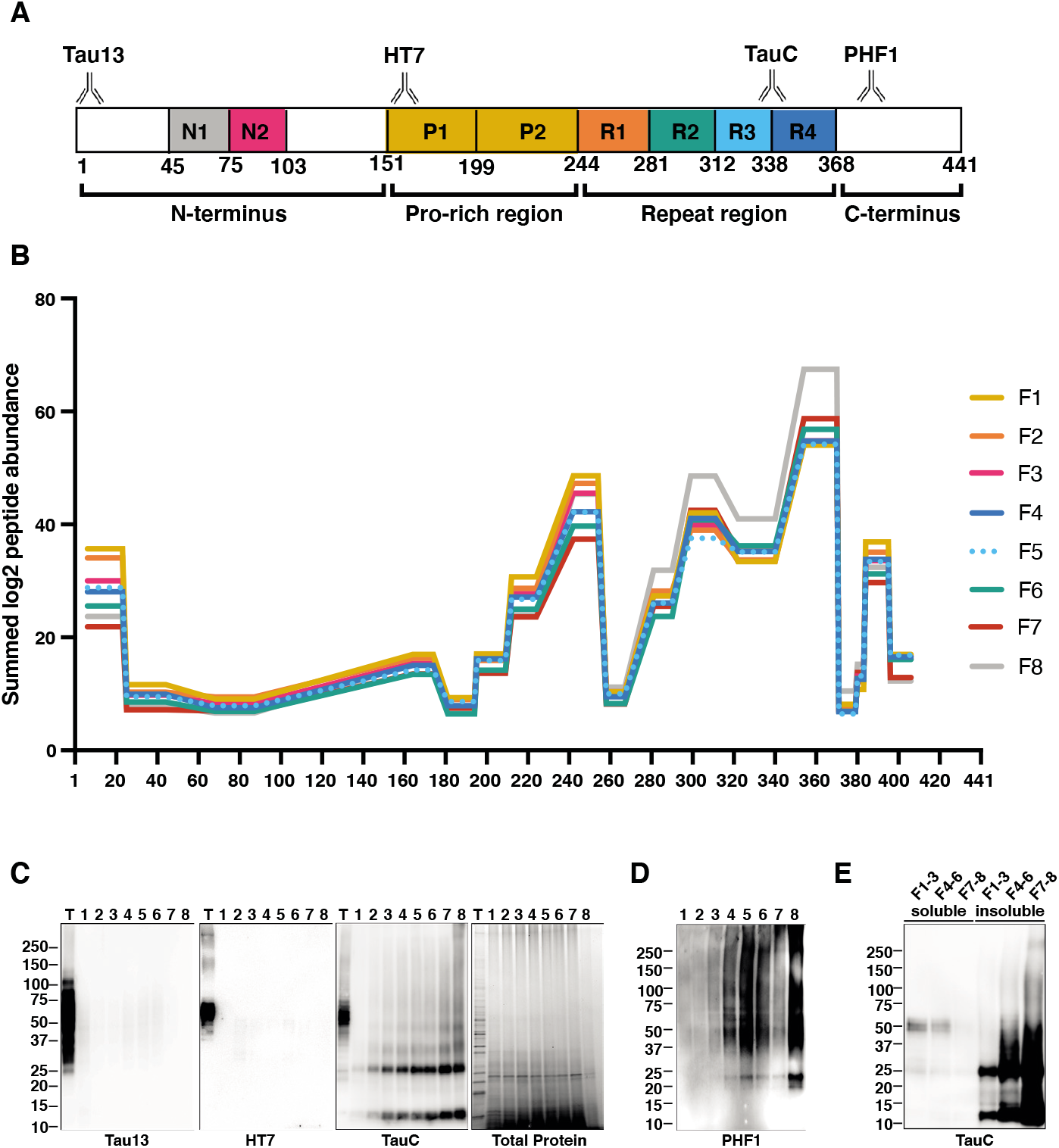
Mass spectrometry and immunoblot analyses of tau protein associated with extracellular vesicles from Alzheimer’s disease patient brain. (A) Schematic of the longest 2N4R tau isoform (441 amino acids) with epitopes shown for antibodies Tau13, HT7, TauC, and PHF1 (phospho-ser396 and -ser404). The tau N-terminal inserts (N1 and N2), proline-rich regions (P1 and P2), and repeats (R1–R4) are shown. (B) Tau peptides identified by quantitative mass spectrometry in EV fractions 1–8. Imputed, normalised, and summed tau peptide label-free quantification intensities are shown (C) Representative immunoblot of total homogenate (T) and EV fractions 1–8 isolated from AD patient brain using antibodies Tau13, HT7 and TauC. Total protein loading control was visualised using the stain-free gel. n=8. (D) Representative immunoblot of EV fractions 1–8 isolated from AD brain using antibody PHF1 (phospho-ser396 and -ser404 tau). n=3. (E) Representative immunoblot of total, sarkosyl-soluble and -insoluble fractions of pooled EV fractions F1-3, F4-6 and F7-8 from AD patient brain using antibody TauC. n=3.

Immunoblotting using the antibody PHF1 (phospho-S396 and -S404 tau) showed that phosphorylated tau was present in fractions 4–6 and 8 (Figure 2D). The 12 kDa tau fragment was not detected by PHF1, consistent with the C-terminal truncation of this fragment. Incubation of the EV fractions with the detergent sarkosyl, followed by differential centrifugation and immunoblotting, demonstrated that the truncated tau was sarkosyl-insoluble and, therefore, assembled (Figure 2E)^1^. To test for contamination by cell-derived sarkosyl-insoluble tau, we spiked brain tissue from individuals without tau pathology with the sarkosyl-insoluble fraction of AD patient brain during tissue dissociation, and proceeded to isolate EVs (Figures S2E and S2F). Sarkosyl-insoluble tau did not co-isolate with the EVs, suggesting that cell-derived sarkosyl-insoluble tau was not contaminating our EV samples. These results show that phosphorylated, assembled truncated tau containing the repeat region is associated with EVs from the brains of AD patients.

### EVs from AD patient brain contain tau PHFs and SFs

We used cryo-EM to image intact EVs from fraction 1, pooled fractions 4–6 and fraction 8 in near-native conditions. EVs in fractions 1 and pooled fractions 4–6 were electron lucent, whereas fraction 8 comprised both electron lucent and electron dense EVs (Figure S3A). A subset of the electron lucent EVs in fraction 8 contained cristae-like membrane structures (Figures S3A and S3B), consistent with their enrichment in mitochondrial proteins. We observed electron lucent EVs in fractions 4–6 that were associated with helical filaments that resembled tau PHFs and SFs by their dimensions and helical crossover spacings (Figure S3A). Similar EVs were not observed in fractions 1 and 8.

We used cryo-ET to reconstruct molecular-resolution tomograms of the EVs associated with tau filaments. This revealed that the filaments were contained within the lumen of the EVs, which had intact limiting membranes (Figures 3A, S4A and S4B). Variable numbers of filaments were observed per EV, ranging from 3 to >50 (Figure 3B). A protease protection assay supported the observation that tau filaments were contained within the lumen of intact EVs. Tau associated with EVs could only be degraded by proteinase K following treatment with detergent to permeabilise EV membranes (Figure 3C).

**Figure 3:**
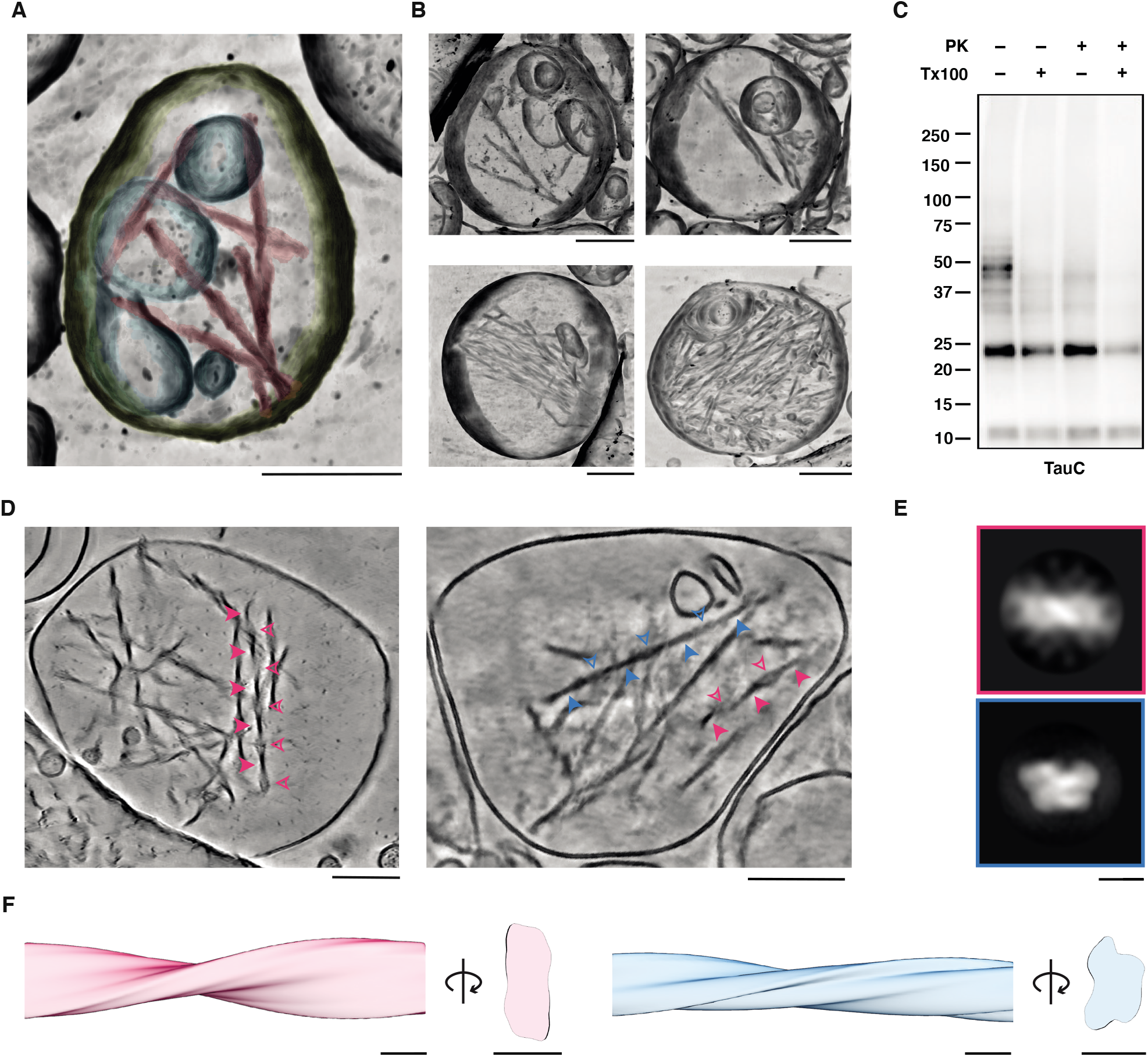
Cryo-ET of extracellular vesicles containing tau PHFs and SFs from Alzheimer’s Disease patient brain. (A) Denoised tomographic volume and segmentation of an EV from AD patient brain depicting the limiting membrane (yellow), intraluminal vesicles (cyan) and tau filaments (magenta). See also Figures S4A and S4B. (B) Denoised tomographic volumes of EVs from AD patient brain containing varying numbers of tau filaments. (C) Representative immunoblot of AD patient brain EVs from fractions 4–6 using antibody TauC with (+) or without (-) 0.1% Triton X-100 and 0.5 ng/µL proteinase K (PK). n=3. (D) Denoised tomographic slices of EVs from AD patient brain containing tau filaments. The arrows point to the minimum (filled arrows) and maximum (unfilled arrows) widths of tau PHFs (magenta arrows) and SFs (blue arrows). (E and F) Subtomogram averaged maps of tau PHFs (magenta) and SFs (blue) in EVs, shown as (E) central slices perpendicular to the helical filament axis and (F) 3D volumes encompassing one helical crossover. See also Figures S4C and S4D. Scale bars, 200 nm in (A, B and D) and 10 nm in (E and F).

Subtomogram averaging of the filaments within EVs yielded two filament types (Figures 3D– 3F, S4C and S4D). The first type resembled tau PHFs, which have a symmetrical, base-to-base packing of the C-shaped protofilaments^1^. The second type resembled tau SFs, in which the protofilaments pack in a non-symmetrical, back-to-back arrangement. This analysis was supported by measurements of single filaments. Tau filaments had average helical crossovers of about 800 nm and cross-sectional dimensions of between 7 and 20 nm (Figures S4E and S4F), in agreement with the dimensions of tau PHFs and SFs^1, 2^. The filaments were between 75 and 500 nm in length (Figure S4G). This is shorter than tau filaments observed within neurofibrillary tangles, which span micrometres^33^. The PHF and SF morphologies made them readily distinguishable from other filaments present within EVs, such as intermediate filaments and actin^34, 35^, which we also observed in our tomograms (Figure S4H).

The EVs containing tau filaments had single limiting membranes and often also contained internal vesicles (Figures 3A, 3B and 3D). Both the EVs and the internal vesicles displayed variable granularity. The EVs were large compared to the majority of EVs, with diameters >300 nm (Figure S4I). These observations were supported by our NTA measurements, which showed that approximately 6% of EVs had diameters >300 nm (Figures S2B–S2D). Our results show that tau PHFs and SFs are contained within the lumen of EVs from AD patient brain.

### Tau filaments associate with one another and are tethered to the EV limiting membrane

Cryo-ET revealed that at least one tau filament per EV was tethered to the luminal side of the EV limiting membrane (Figure 4A). The remaining filaments made lateral contacts with the tethered filament and/or with one another, resulting in a polarised filament distribution (Figures 3A, 3B and 3D). Tethering to the EV limiting membrane occurred exclusively at filament ends and was mediated by flexible elongated densities that often appeared to traverse the limiting membrane (Figure 4A). Fractionation of EVs into surface proteins, luminal proteins and membrane-associated proteins^36^ supported the observation that the filaments were tethered to the limiting membrane. Tau was retained in the membrane fraction following treatment of EVs with 0.1 M sodium carbonate pH 11, demonstrating membrane interaction (Figure 4B).

**Figure 4:**
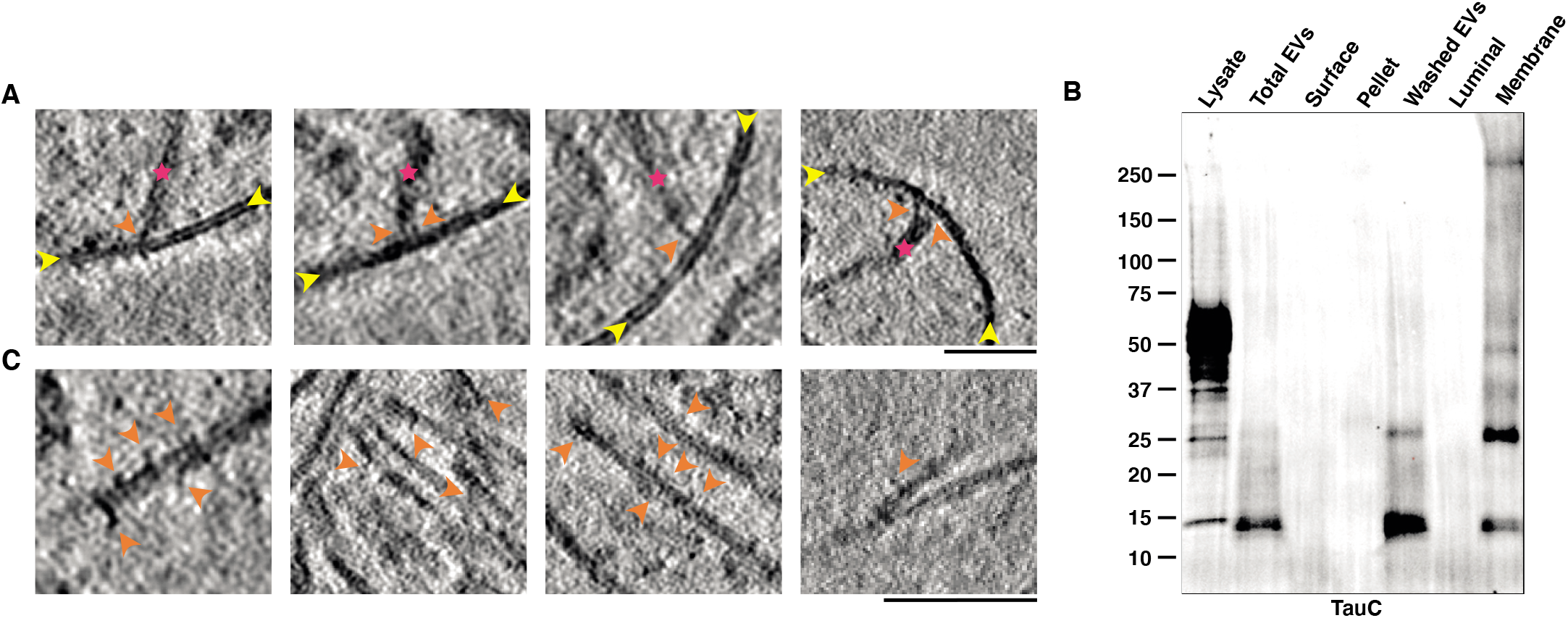
Tau filaments are tethered to the limiting membrane of extracellular vesicles from Alzheimer’s disease patient brain and are decorated by additional densities. (A) Deconvolved tomographic slices of EVs from AD patient brain showing tethering of tau filaments to the luminal side of the limiting membrane. The magenta asterisks indicate filaments, the yellow arrows point to the limiting membranes of EVs, and the purple arrows point to densities tethering the filaments to the luminal side of the limiting membrane. (B) Representative immunoblot using antibody TauC of total EVs; proteins liberated from the EV surface following DTT wash (Surface); sedimented material (Pellet; not associated with membranes); DTT-washed EVs (Washed EVs); luminal proteins following resuspension of DTT-washed EVs in hypotonic buffer (Luminal); and the sodium carbonate-treated membrane-associated fraction (Membrane). n=3. (C) Deconvolved tomographic slices of EVs from AD patient brain showing additional densities decorating the lengths of tau filaments. Purple arrows point to additional densities. Scale bars, 200 nm in (A and C).

Occasionally, the sides of filaments also made contact with internal vesicles, in the absence of linking densities (Figures 3A and 3B). We also observed globular densities that decorated the sides of the filaments (Figure 4C). Some of these densities followed the helical symmetry of the filaments, suggesting that they associated with specific tau sequences or structural motifs. These results show that tau filaments within EVs in AD patient brain associate with one another and are tethered to the limiting membrane.

### Tau PHFs within EVs from AD patient brain contain additional molecules

We used single-particle cryo-EM to determine the structure of tau PHFs extracted from EVs isolated from the brains of two AD patients (Figures 5A, 5B and S5A–D). This revealed that the filaments had structures similar to PHFs from total brain homogenates of AD patient brain ^1, 2^. The two C-shaped protofilaments had a combined cross-β/β-helix structure formed by residues S305–R379 of 3R and 4R tau isoforms. They were related by helical 2-1 screw symmetry, with a protofilament interface formed by residues K331–E338.

**Figure 5:**
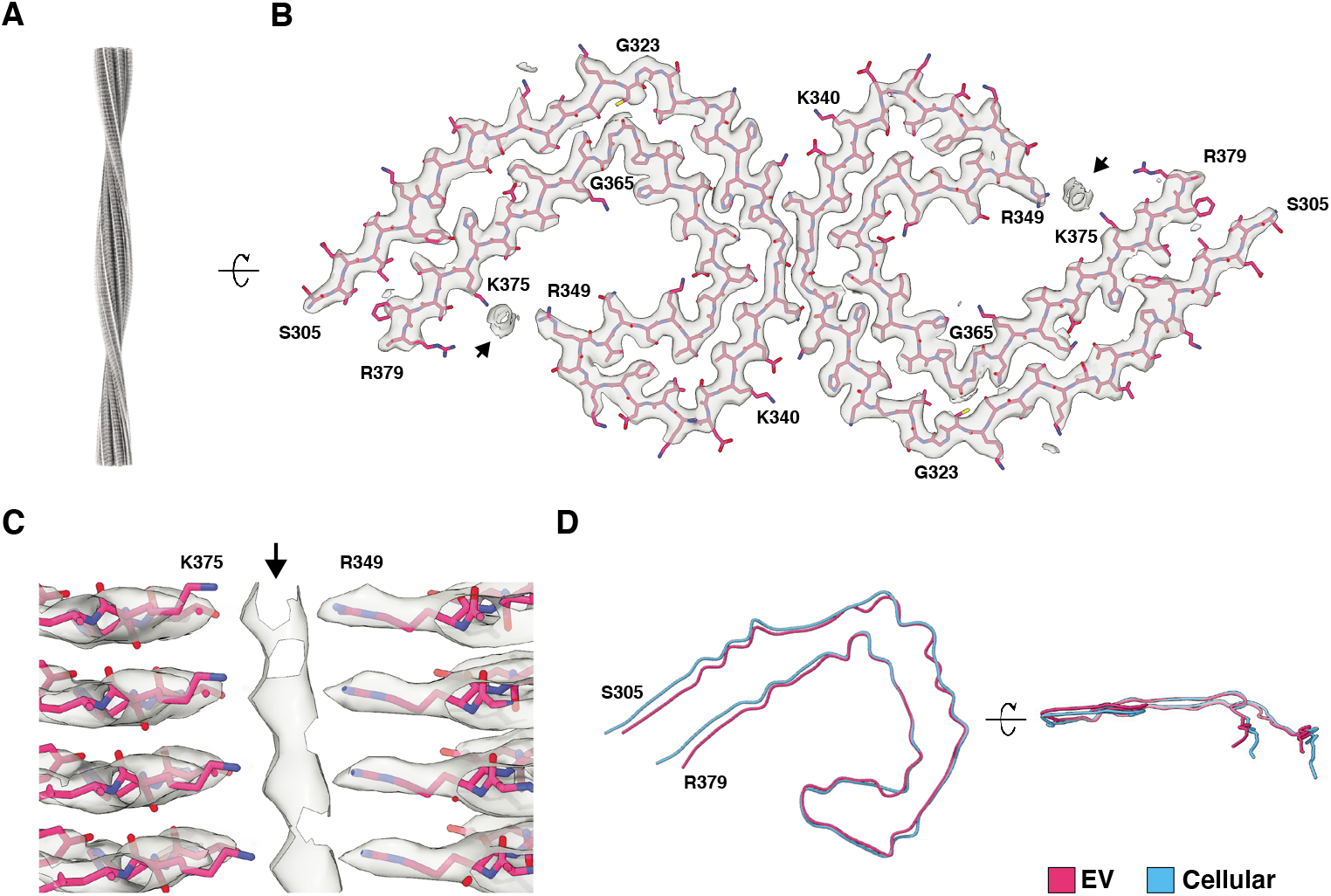
Cryo-EM structure of extracellular vesicle-derived tau PHFs from Alzheimer’s disease patient brain. (A) Cryo-EM map of tau PHFs from AD patient brain EVs, viewed aligned to the helical axis. See also Figures S5A–E. (B) Cryo-EM map (transparent grey) and atomic model (magenta) of tau PHFs from AD patient brain EVs, shown for a single tau molecule per protofilament perpendicular to the helical axis. Additional densities coordinated to the side chains of R349 and K375 are indicated with arrows. (C) Magnified view of the cryo-EM map (transparent grey) and atomic model (magenta) showing the additional density, indicated with an arrow, coordinated to the side chains of R349 and K375. Four tau molecules are depicted aligned to the helical axis. (D) Overlay of the tau filament folds from EVs (magenta) and the cellular fraction (cyan) from AD patient brain. See also Figure S5F.

Unlike PHFs isolated from total brain homogenates and PHFs reconstituted *in vitro* from a fragment of tau comprising residues 297–391^1, 2, 37^, the PHFs from EVs contained additional densities between the positively-charged side chains of R349 and K375, revealing the presence of anionic molecules (Figures 5B, 5C and S5E). Their identity was not resolved, possibly because they do not share the helical symmetry of the tau molecules. These molecules were accompanied by a more compacted C-shaped fold than that of PHFs from total brain homogenates and of PHFs reconstituted *in vitro* (Figures 5D and S5F). Compaction was mediated by hinging of the mainchain at glycine residues 323 and 366 in the cross-β region, and at K340 and I354 in the β-helix region.

We also determined the structure of PHFs extracted from the cellular brain fraction from one of the AD patients (Figures 5D and S5). This was identical to those determined from total homogenates and reconstituted *in vitro*^1, 2, 37^. We conclude that PHFs within brain EVs from AD patients contain additional anionic molecules compared to those within neurofibrillary tangles.

## DISCUSSION

The association of assembled tau with EVs in the central nervous system of patients with AD has been linked to its clearance and prion-like propagation^20–23^. However, our understanding of the molecular mechanisms underlying EV-mediated secretion of assembled tau is limited, because the molecular species of assembled tau; the identities of the EVs that associate with assembled tau; and how tau associates with EVs have not been determined.

We carried out a comprehensive density-based profiling of brain EVs from AD patients. This revealed the enrichment of proteins from distinct intracellular compartments of EV biogenesis in different EV fractions, including from the plasma membrane, endo-lysosomes and mitochondria. We found that EVs containing tau filaments were enriched in endo-lysosomal proteins. Association of lysosomal substrates with EVs has been observed as a consequence of lysosomal impairment^38–40^, including assembled protein substrates^41, 42^. This suggests that tau filaments may be secreted in EVs as a result of impaired lysosomal degradation. Tau filaments are targeted to lysosomes in cell and *in vivo* models^43^. Lysosomal impairment has been described in AD, and multiple genetic risk factors for AD are implicated in lysosomal dysfunction^43^.

It was not known which molecular species and structures of assembled tau associate with EVs. We showed that PHFs and SFs composed of tau truncated at the N- and C-termini are found within the lumen of EVs in AD patient brain. Truncation is compatible with the cryo-EM structure of the C-shaped protofilament fold, which encompasses residues 305–379. The enrichment of peptides from the extreme N-terminus of tau may reflect the discontinuous conformational MC1 epitope of PHFs and SFs, which comprises tau residues 7–9 and 313– 312^44^. Filaments of truncated tau have previously been observed in total homogenates of AD patient brain and have been shown to seed tau assembly^45, 46^. Our results suggest that tau proteolysis could occur within the protease-rich environment of EVs, or in the endo-lysosomal pathway prior to EV biogenesis. Our finding that tau is truncated and that the residues S305– R379 are buried in the ordered filament core will inform the development of tools that target assembled EV-associated tau for therapeutic and biomarker applications.

Tau PHFs and SFs within EVs were short compared to those in neurofibrillary tangles. Short tau filaments generated by homogenisation and fractionation of human or transgenic mouse brain have been shown to have the greatest ability to seed tau assembly^11, 47–49^. However, it was not known if short tau filaments existed *in situ* in human brain. Our results suggest that short tau filaments have the potential to act as seeds for prion-like propagation in the human brain.

We observed additional anionic molecules in PHFs from EVs compared to PHFs from the cellular fraction, demonstrating that tau filament composition can differ between cellular compartments. The anionic molecules may be derived from the EV lumen, or from environments upstream of EV biogenesis. We also observed additional globular densities decorating the sides of the filaments by cryo-ET, which are not observed on filaments extracted from human brain and may be lost during extraction^1, 2^. Co-immunoprecipitation experiments have shown that a large number of proteins interact with tau filaments in human brain^49^. The identities of the molecules that interact with tau filaments in EVs, as well as if they influence filament formation and EV-mediated secretion, remain to be determined.

How assembled tau associates with EVs was unknown. Our results reveal that tau filaments are contained within the lumen of EVs. This finding has important implications for therapeutic approaches targeting extracellular tau, which will be inaccessible to antibodies and membrane-impermeable compounds. Our observation that densities tether tau filaments to the EV limiting membrane suggests that the filaments may be actively sorted into EVs. Tethering may require adaptor molecules, such as Bin1^21^, or may be mediated by direct polypeptide-anchoring to membranes^50^. That filaments were uniquely tethered at their ends suggests that sequences or structural motifs exposed at the filament ends are important for membrane association. Targeting of the tethering mechanism may represent a strategy to modulate EV-mediated secretion of tau filaments.

We have shown that PHFs and SFs of truncated tau are tethered within EVs enriched in endo- lysosomal proteins in AD patient brain. This work paves the way for further *in situ* cryo-ET studies of neurodegenerative disease-associated protein assemblies in human brain. Our results will guide mechanistic studies into the secretion of tau filaments within EVs, as well as inform strategies to target extracellular tau.

## ACKNOWLEDGEMENTS

We thank the individuals and their families for donating brain tissue; Columbia University New York Brain Bank and the University of Miami Brain Endowment Bank for access to tissue resources and pathological expertise; Peter Davies for his generous donation of tau antibodies; Alister Burt and Tom Dendooven for help with analysing the cryo-ET data; Keitaro Yamashita and Garib Murshudov for help with cryo-EM model refinements in Servalcat and REFMAC5; staff at the MRC Laboratory of Molecular Biology Electron Microscopy Facility for access to and support with cryo-EM; staff at the MRC Laboratory of Molecular Biology Scientific Computing Facility for access to and support with computing; the European Synchrotron Radiation Facility (ESRF) for provision of beam time on CM01; and Gregory Effantin and Diana Arseni for support with cryo-EM at the ESRF. This work was supported by the National Institutes of Health (AG05717 and AG056732 to E.L.; and AG063521 and AG056151 to K.E.D.); the UK Dementia Research Institute, which receives funding from DRI Ltd., funded by the Medical Research Council, as part of United Kingdom Research and Innovation (also known as UK Research and Innovation), the Alzheimer’s Society and Alzheimer’s Research UK to K.E.D; the BrightFocus Foundation (A2017393S to K.E.D.); the Medical Research Council, as part of United Kingdom Research and Innovation (also known as UK Research and Innovation) (MC_UP_1201/25 to B.R.-F.); and Alzheimer’s Research UK (ARUK-RS2019-001 to B.R.-F.).

## AUTHOR CONTRIBUTIONS

S.L.F. performed EV isolations and EV biochemistry; S.L.F. and T.S.B. performed tau filament extraction; T.S.B. performed cryo-ET and subtomogram averaging; T.S.B. and B.R.-F. performed single-particle cryo-EM; S.L.F. and E.T. performed proteomic and NTA bioinformatic analyses; S.L.F. and P.M.C. performed NTA; S.L.F performed HEK biosensor cell seeding; A.S. and G.C. performed mouse stereotaxic injections; M.S.F. performed mouse brain immunostaining and analysis; S.B, E.F, and E.T assisted in EV isolation and analysis; D.P.O. and R.F performed LC MS-MS; D.P.O., B.G. and P.G. consulted on LC MS-MS data analysis; C.M. performed initial EV isolations from human tissue; P.D.A. and E.L. consulted on EVs; E.L., K.E.D. and B.R.-F. acquired funding; K.E.D. and B.R.-F. supervised the study; S.L.F., T.S.B., K.E.D. and B.R.-F wrote the manuscript.

---

**Table S1:** Human Brain Tissue Samples.

| Donor | Sex | Age | PMI cold | PMI frozen | ABC score | aSyn | TDP43 | Diagnosis | LC MS-MS region | Cryo-EM region | Cryo-ET region |
| --- | --- | --- | --- | --- | --- | --- | --- | --- | --- | --- | --- |
| **AD cases** |  |  |  |  |  |  |  |  |  |  |  |
| A | M | 67 | 0:00 | 14:55 | A3, B3, C3 | - | - | ADNP | HH | - | - |
| B | M | 89+ | 7:40 | 19:21 | A3, B3, C3 | - | - | ADNP | TC | - | TC, H |
| C | F | 68 | 3:30 | 9:20 | A3, B3, C3 | - | - | ADNP | FC | H, TC, HH, A | - |
| D | M | 73 | 0:00 | 23:00 | A3, B3, C3 | - | - | ADNP | H | - | - |
| E | M | 89 | 0:30 | 7:17 | A2, B3, C3 | - | - | ADNP | TC | - | - |
| F | M | 88 | 0:00 | 6:00 | A3, B3, C3 | - | - | ADNP | H | - | - |
| G | F | 89+ | 2:15 | 6:16 | A3, B3, C3 | - | - | ADNP | TC | - | - |
| H | F | 89+ | 1:00 | 19:15 | A3, B3, C3 | - | * | ADNP | HH | - | - |
| I | M | 83 | 0:00 | 17:16 | A3, B3, C3 | - | - | ADNP | - | H, TC, HH, A | - |
| **Low tau pathology cases** |  |  |  |  |  |  |  |  |  |  |  |
| J | F | 47 | N/A | 12:00 | - | - | - | NP normal | - | - | - |
| K | M | 54 | N/A | 7:30 | - | - | - | NP normal | - | - | - |

PMI cold: post-mortem interval – time to cold (h:min); PMI frozen: post-mortem interval – time to frozen (h:min). All AD cases were diagnosed on autopsy as ADNC (Alzheimer's Disease Neuropathologic Changes), were classified as Braak stage VI, with high likelihood that dementia was due to AD. All AD cases were categorized as A3 (Thal phase for A $\beta$ plaques 4 or 5), B3 (Braak stage for tau neurofibrillary tangles V or VI), C3 (CERAD neuritic plaque score of 'Frequent') except case E (A2). aMI: acute myocardial infarction; TC: temporal cortex; HH: head of hippocampus; H: hippocampus, FC: frontal cortex; A: amygdala; PVC: primary visual cortex; Cryo-EM: Single-particle cryo-electron microscopy; Cryo-ET: cryo-electron tomography; aSyn: concomitant alpha-synuclein pathology; TDP43: concomitant TDP-43 pathology; NP normal: neuropathologically normal. \*Rare neurons in temporal regions with ubiquitinated TDP-43 labelling. All human AD specimens were fresh frozen and sourced from the New York Brain Bank (NYBB) at Columbia University (Alzheimer's Disease Research Center). Low tau pathology cases were sourced from the University of Miami Brain Endowment Bank.

**Table S2:** Cryo-EM data collection, refinement and validation statistics.

|  | **EV 1**<br>PDB: 8BGS<br>EMDB: 16035<br>EMPIAR: 11300 | **EV 2** | **Cell**<br>PDB: 8BGV<br>EMDB: 16039<br>EMPIAR: 11301 |
| --- | --- | --- | --- |
| **Data collection and processing** |  |  |  |
| Voltage (keV) | 300 | 300 | 300 |
| Electron source | XFEG | XFEG | XFEG |
| Detector | K2 | K3 | K3 |
| Energy filter slit width (eV) | 20 | 20 | 20 |
| Electron exposure (e<sup>-</sup>/Å<sup>2</sup>) | 38.7 | 40 | 40 |
| Defocus range (μm) | -2.2 to -1.4 | -2.4 to -1.0 | -2.4 to -1.0 |
| Pixel size (Å) | 1.05 | 0.93 | 0.834 |
| Symmetry imposed | C1 | C1 | C1 |
| Initial particle images (no.) | 287,758 | 87,847 | 98,918 |
| Final particle images (no.) | 18,041 | 19,059 | 9,347 |
| Helical twist (°) | 174.439 | 174.443 | 174.454 |
| Helical rise (Å) | 2.354 | 2.417 | 2.413 |
| Map resolution (Å) | 3.16 | 3.40 | 3.27 |
| FSC threshold | 0.143 | 0.143 | 0.143 |
| Map sharpening *B* factor (Å<sup>2</sup>) | -39.35 | -61.15 | -54.79 |
| **Refinement** |  |  |  |
| Model resolution (Å) | 3.24 | — | 3.31 |
| FSC threshold | 0.5 | — | 0.5 |
| Model composition |  |  |  |
| Non-hydrogen atoms | 574 | — | 587 |
| Protein residues | 75 | — | 77 |
| Ligands | 0 | — | 0 |
| *B* factors (Å<sup>2</sup>) |  |  |  |
| Protein | 69.32 | — | 66.68 |
| Ligand | — | — | — |
| R.m.s. deviations |  |  |  |
| Bond lengths (Å) | 0.0086 | — | 0.0084 |
| Bond angles (°) | 1.78 | — | 1.79 |
| Validation |  |  |  |
| MolProbity score | 1.32 | — | 1.19 |
| Clashscore | 1.46 | — | 0.84 |
| Poor rotamers (%) | 0 | — | 0 |
| Ramachandran plot |  |  |  |
| Favoured (%) | 93.15 | — | 93.33 |
| Allowed (%) | 100 | — | 100 |
| Outliers (%) | 0 | — | 0 |

## METHODS

### Isolation of extracellular vesicles from post-mortem human brain tissue

EVs were isolated essentially as described^32^, with minor alterations to achieve a gentler tissue dissociation suitable for post-mortem human brain tissue. Briefly, grey matter from flash-frozen post-mortem human brain tissue (Table S1) was dissected on ice and minced using two razor blades in a few drops of pre-warmed (37°C) Hibernate A medium containing 20 units/mL papain (Worthington LK003178). Minced tissue was incubated in 3.125 mL pre-warmed papain solution for 10 minutes at 37°C, with three gentle tube inversions after 5 min. Papain digestion was stopped by the addition of 6.5 mL ice-cold Hibernate A medium containing protease inhibitors (5 μg/mL leupeptin, 5 μg/mL antipain dihydrochloride, 5 μg/mL pepstatin A, 1 mM phenylmethanesulfonyl fluoride, and 1 μM E64), and the dissociated tissue was passed slowly through a 10 mL pipette until smooth (6-8 times). The dissociated tissue was then centrifuged at 300 x g for 10 min at 4°C to pellet cells. The supernatant, containing the extracellular components, was passed through a 40 μm cell strainer (Corning 352340), followed by a low-protein binding 0.2 μm syringe filter (Corning 431219). The filtrate was centrifuged at 2,000 x g for 10 min at 4°C and the supernatant was made to 60 mL with ice-cold PBS, and further centrifuged at 10,000 x g for 30 min at 4°C. The supernatant was transferred to fresh tubes and centrifuged at 100,000 x g for 70 min at 4°C using a Ti45 rotor (Beckman). The 100,000 x g pellets were resuspended in 1 mL ice-cold PBS and brought to 60 mL using PBS, before another round of centrifugation at 100,000 x g for 70 min at 4°C. The supernatants were discarded and the pellets (containing EVs) were resuspended in 1.5 mL 60 mM Tris-HCl, pH 7.4, containing 0.25 M sucrose and 40% Optiprep (OP) and placed in ultra-clear centrifuge tubes (Beckman, 344059). Discontinuous OP gradients were prepared by layering 2 mL of 60 mM Tris-HCl, pH 7.4, containing 0.25 M sucrose and 20%, 15%, 13%, 11%, 9%, 7%, and 5% OP, respectively. Additional 60 mM Tris-HCl, pH 7.4, containing 0.25 M sucrose and 5% OP was then added until 1-3 mm from the top of the tube. The prepared gradients were centrifuged at 200,000 x g for 16 h at 4°C in an SW41Ti rotor (Beckman), with no braking during deceleration. Eight fractions encompassing each OP concentration were collected into 30 mL thickwall polycarbonate tubes (Beckman 355631) as described in Figure S1 (F1 = 5% OP; F2 = 7% OP; F3 = 9% OP; F4 = 11% OP; F5 = 13% OP; F6 = 15% OP; F7

= 20% OP; F8 = interface between the 20/40% OP layers). Fractions were diluted with 15 mL of cold PBS and centrifuged at 100,000 x g for 70 min at 4°C using a 70.1Ti rotor (Beckman). The supernatants were discarded and the pellets (containing EVs) were resuspended in 50 μL PBS. EV suspensions were used immediately for cryo-ET and for proteinase K protection assays, or aliquoted and frozen at -80°C for additional biochemical analyses, including prior to sarkosyl extraction for cryo-EM.

### Immunoblotting

EV suspensions in PBS were diluted 1:1 in 2X RIPA (100 mM Tris-HCL, pH 7.4 300 mM NaCl, 2% NP-40, 1% sodium deoxycholate, 0.2% SDS). Total protein concentration was determined using a bicinchoninic acid (BCA) assay. EV suspensions were normalised to total protein concentration, prepared with 4X Laemmli sample buffer (BioRad; 1610747) containing 50 mM dithiothreitol (DTT), and heated at 95°C for 5 min. Samples were resolved using 4-20% tris-glycine extended (TGX) stain-free gels (BioRad). Proteins were transferred to low-fluorescence PVDF membranes (BioRad) and visualised using UV light in a Chemi-Doc MP imager (BioRad). Membranes were subsequently blocked with 5% milk powder in tris-buffered saline (20 mM Tris-HCl pH 7.4, 150 mM NaCl) containing 0.02% Tween-20 (TBST) for 1 h at 21°C. Primary antibodies were diluted in Superblock TBS blocking buffer (Thermo Scientific) and incubated overnight with the membranes. Membranes were then washed twice in TBST and twice in TBST containing 5% milk powder, with each wash lasting 10 min. Secondary HRP-conjugated antibodies were diluted in TBST containing 5% milk powder and incubated with the membranes for 1 h at 21°C. Membranes were then washed four times in TBST for 10 min each wash and labelled proteins were visualised using Bio-Rad Clarity Max chemiluminescent substrate in a Chemi-Doc MP imager. The total protein signal visualised using UV light was used to normalise the labelled protein signal using the Image Lab software package Version 6.1, build 7 (BioRad). Primary antibodies were as follows: annexin A2 (Cell Signaling D11G2, 1/1000), flotillin 1 (BD Transduction 610821, 1/1000), CD81 (Cell Signaling D3N2D, 1/1000), LAMP2 (Santa Cruz sc-18822 clone H4B4, 1/500), VDAC (Cell Signaling 48665, 1/1000), lamin A/C (Santa Cruz sc-376248, 1/1000), Tau13 (BioVision 3453-100, 1/1000), HT7 (Invitrogen MN1000, 1/1000), TauC (DAKO A0024, 1/5000), PHF1 (mouse monoclonal; kind gift from Peter Davies, 1/250).

### Proteomic sample preparation and LC-MS/MS analysis

#### Reduction, alkylation, S-TRAP clean-up and protease digestion

EV fractions were resuspended in TBS (20 mM Tris-HCL, pH 7.4, 150 mM NaCl) containing 5% SDS, and proteins were digested using S-TRAP Micro spin columns, according to the manufacturer’s protocol (Profiti). Approximately 50 μL of starting material was used for each experiment, corresponding to ∼50 μg of total protein. Samples were reduced by the addition of 20 mM DTT and incubated for 30 min at 21°C. Samples were alkylated by the addition of 40 mM iodoacetamide for 30 min at 21°C, protected from light. Phosphoric acid was then added to each sample to 1.2 %, followed by vortexing. Samples were diluted 6.6-fold in S-TRAP Bind and Wash buffer composed of 90% methanol containing 100 mM tetraethylammonium bromide (TEAB), pH 8, and loaded into an S-TRAP Micro spin column by centrifugation at 4,000 x g at 21°C until all of the solution passed through the column. Each spin column was subsequently washed with 150 μL of S-TRAP Bind and Wash buffer and centrifuged at 4,000 x g for 30 s at 21°C. The flow-through was discarded. This wash step was repeated 3 more times for a total of 4 washes. Prior to digestion, spin columns were transferred to clean collection tubes. Porcine trypsin (Promega) in 50 mM TEAB, pH 8, was added to each spin column to a final enzyme:substrate ratio of 1:10, followed by incubation for 2 h at 47°C without agitation. Digested peptides were eluted from the spin columns by the addition of 40 μL of S-TRAP Elution Buffer A composed of 50 mM TEAB, pH 8, and centrifugation at 4,000 x g for 30 s at 21°C. A second elution was carried out by the addition of equal volumes of S-TRAP Elution Buffer B composed of 0.5% trifluoroacetic acid (TFA) in water and S-TRAP Elution Buffer C composed of 0.5% TFA in 50% acetonitrile. The eluates were combined, dried using a speedvac and resuspended in 0.1% formic acid (FA) for subsequent mass spectrometry (MS) analysis.

#### Evosep/timsTOF LC-MS/MS

A high throughput LC-MS/MS workflow was implemented as described in^51^. EvoTips (Evosep) were activated by soaking in propanol, then washed with 20 μL of 0.1% FA in acetonitrile (solvent B) with centrifugation at 700 x g for 60 s. Tips were conditioned by soaking in propanol until the C18 material appeared pale white, centrifuged at 700 x g for 60 s, and equilibrated with 20 μL of 0.1 % FA in water (solvent A). To prevent drying out, samples were loaded into tips while equilibrating in solvent A. Peptides were bound to the C18 material by centrifugation at 700 x g for 60 s. Tips were subsequently washed with 20 μL solvent A and centrifugation at 700 x g for 60 s. Following this, 100 μL of solvent A was added to the tips, which were then centrifuged at 700 x g for 10 s. LC-MS/MS analysis was performed immediately. Peptides were analysed using an Evosep One (Evosep) coupled to a timsTOF Pro mass spectrometer (Bruker), with separation of peptides on a C18 column (150 μm x 80 mm) packed with 1.5 μm beads (PepSep). A gradient length of 21 min at a flow rate of 1 μL/min was used. The timsTOF Pro mass spectrometer was operated in a parallel accumulation, serial fragmentation (PASEF) mode. Trapped ion mobility spectrometry (TIMS) ion accumulation and ramp times were set to 100 ms and mass spectra were recorded from *m/z* 100 – 1700. diaPASEF used 8 diaPASEF scans per TIMS-MS scan, giving a duty cycle of ∼1 s. The ion mobility range was set to 0.85 – 1.3 Vs/cm^2^. Each mass window isolated was *m/z* 25 wide, ranging from *m/z* 475-1000, and with an ion mobility-dependent collision energy that increased linearly from 27 eV to 45 eV between 0.85 – 1.3 Vs/cm^2^.

#### Data searching

Data-Independent Acquisition (DIA) data was searched using default settings in DIA-NN software (version 1.8). Methionine oxidation was set as a variable modification and cysteine carbamidomethylation set as a fixed modification. The search allowed for one missed trypsin cleavage site. Samples were searched against the *Homo sapiens* Uniprot database (retrieved in Feb 2021) and quantified by label-free quantitation (LFQ).

#### MS Data analysis

Raw protein intensities were analysed in R 4.2.1^52^ with all code documented within R Markdown^53^ at https://github.com/duff-lab-team/AD-EV-characterisation. Reproducibility and package versioning is provided through Singularity/Docker^54^. The following pipeline was used for both protein and peptide LFQ intensities. Intensity data was imported and handled using the Differential Enrichment analysis of Proteomics data package^55^. A total of 6105 unique proteins were identified across all the fraction groups. Three F8 samples (A, F, and E) were removed from the analysis as significantly less material was isolated from those donors. Post filtering (described in the flowchart below), 6054 proteins were retained for the imputation pipeline:

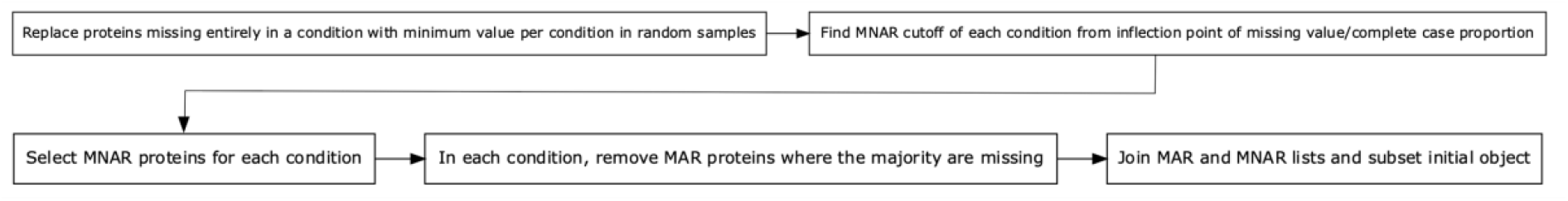

Missing data (Figure S2B) were handled using a hybrid missing-not-at-random (MNAR) and missing-at-random (MAR) imputation strategy designed to distinguish between proteins that were completely absent from a particular fraction (likely MNAR), from proteins with values missing only from a small number of donors or fractions (likely MAR). This hybrid strategy maximises the inclusion of differentially expressed proteins (DEPs) across the fractions and avoids exclusion of biologically relevant protein identities (eg. proteins expressed selectively in one or two fraction groups)^56^. This strategy led to in a 43% increase in the number of differentially expressed proteins than if the analysis was restricted to proteins without missing values in any of the fractions.

#### Design and implementation of the hybrid imputation strategy

To demonstrate the effectiveness of the hybrid imputation strategy and aid in the selection of MAR vs. MNAR missing values, a simulated dataset was produced from a matrix of 6105 complete case log2 intensity values based on the mean and standard deviation of proteins in the real dataset^56^. To model the characteristics of the real dataset, 3000 DEPs, and missing values of both MAR and MNAR-type were manually assigned to the simulation. The relationship between true missing values across different mean intensities with MAR alone (Figure S3A) versus MNAR + MAR (Figure S3B) was investigated to define a cut-off for MNAR missing values. In the simulation, it was determined that setting the MNAR cut-off at the bottom of the steep decline in true missing values (log2 intensity value of 12) (Figure S3B). This maximised the number of true DEPs captured, the accuracy of their detection and the average adjusted F p-value of an ANOVA across all three fraction groups (Figure S4A). Imputation of the simulated dataset identified 92.97% of the true DEPs with similar accuracy to using the complete cases, whilst only 69.07% and 24.03% of true DEPs were captured in the unimputed and complete cases (Figure S4B,C). To apply this strategy to the real dataset, MNAR log2 intensity cut-off values were manually determined for each fraction using the simulation curve as a guide (Figure S3B): F1, 14.9; F2, 14.7; F3, 14.9; F4, 14.3; F5, 14.6; F6, 14.5; F7, 15.3; F8, 15.2. In the real dataset, 827 more proteins were detected as DEPs following imputation (Figure S4D). Data were normalised using a variance stabilising transformation (VSN), and principal component analysis was performed visualised using the R package ggplot2 ^57^. The 68% confidence interval was used to draw ellipses surrounding the points for each donor (Figure 1B).

#### Differential expression and gene ontology pathway analysis

Differential expression analysis was performed using moderated F and t-statistics from the R package limma^58, 59^, with visualisation aided by the R packages volcano3D^60^ and ComplexHeatmap^61^. Pairwise contrasts were tested across the three fraction groups using the Benjamini and Hochberg multiple testing correction factor. Proteins were deemed significant at adjusted-p values < 0.05, without a log fold-change cut-off (Data S1). Gene ontology analysis (Fig 1C) was performed using the ssGSEA method from the GSVA R package^62^. After the ssGSEA transformation, the same limma testing procedure was applied to the new matrix, with a more stringent threshold of adjusted-p < 5x10^-4^. To reduce multiple testing burden, data were filtered such that only gene sets with between 5-50 genes detected were used. Gene sets with more than 25% overlapping genes, and gene sets with directionality modifiers as “up/downregulation of” etc were discarded. Relevant terms with the highest F p-values across the fraction groups were selected for display in Figure 1C (Data S2).

### NTA Particle Analysis

At isolation, fresh EVs were partitioned into 3 μL aliquots for particle counting and frozen at - 80°C. Samples were thawed only once prior to NTA analysis (ZetaView PMX-420 Quatt, Particle Metrix, Germany), and immediately diluted in particle-free PBS to the optimal reading range of the instrument (100-150 particles). Samples were diluted in PBS (between 1/100,000 and 1/5,000) and analysed at 22°C in scatter mode at the following settings: sensitivity = 80, shutter = 100, frame rate = 30, trace length = 15, bin size = 5 nm, positions per single reading = 11. Code and R Markdown sheets describing the data analysis are deposited at: https://github.com/duff-lab-team/AD-EV-characterisation.

### Seeded tau aggregation in Tau RD P301S FRET Biosensor cells

Tau RD P301S FRET Biosensor cells (ATCC CRL-3275) were plated at 30,000 cells per well in 96-well flat-bottomed plates in 130 μL Dulbecco’s Modified Eagle Medium containing 10% FBS and 5% PenStrep. The cells were allowed to settle undisturbed for 15 mins at RT prior to being moved to the incubator and grown at 37°C with 5% CO_2_. The following day, seed mixes were made in 20 μL volumes containing 15 μL protein dilution (3 μg EVs in PBS diluted in a final volume of 15 μL Optimem media) and 5 μL Lipofectamine dilution (1 μL Lipofectamine 2000 plus 4 μL OptiMEM media). Seeding reactions were incubated for 30 mins in the cell culture incubator, before being added to each well of cells 24 h post-plating (cells were ∼65- 70% confluent). The seeded cells were maintained undisturbed in the incubator for 48 h prior to harvesting for FRET measurements.

At time of harvest, the cell media was removed from each well using a vacuum manifold for 96-well plates. Trypsin-EDTA (50 μL) was added to each well, and the plate was incubated for 2-3 minutes at 37°C. Serum-containing media (150 μL) was added to each well, cells were triturated 10X, and moved into wells of a fresh v-bottomed 96-well plate. Cells were pelleted at 600 x g for 5 mins and resuspended in 150 μL cold 4% paraformaldehyde (PFA) in PBS by triturating 10X. Cells were left to fix for 10 mins, and then re-pelleted at 600 x g for 5 mins. The cells were resuspended in 200 μL PBS containing 1 mM EDTA. Flow cytometry analysis of FRET signals was performed using a BD LSR Fortessa flow cytometer equipped with a high throughput sampler using BV510 and BV421 BD filter sets, as previously described^63^. Samples were run on the “low” flow setting (<1000 events/s) and 50,000 events were captured per well, with 3 technical replicates for each condition. Integrated FRET densities (% FRET positive cells per gated cell population x median fluorescence intensity of the BV510 signal) were obtained using FCS Express 7 Research software (De Novo Software).

### Seeded tau aggregation in PS19 mice

All experiments were performed in accordance with protocols approved by the Institutional Animal Care and Use Committee at Columbia University (NYC). Three 6-month-old male PS19 mice were injected with pooled AD EVs (F1-F8) in the left hemisphere, and pooled control EVs (F1-F8) in the right hemisphere. Mice were immobilized in a stereotaxic frame (David Kopf Instruments) under continuous isofluorane administration and stereotaxic injections were made under aseptic conditions using a Hamilton syringe into the hippocampal hilus (AP, − 2.5 mm; ML, +/− 2 mm; and DV, −1.8 mm). Each injection contained 10 μg of total EV protein in a volume of 1.5 μL. All animals were carefully monitored and administered analgesics both during and after surgery. Mice were sacrificed 2 months post-injection and perfused intracardially with PBS and 10% formalin. Brains were postfixed overnight in 10% formalin, and cryoprotected in 30% sucrose for 1 week prior to TissueTek embedding. Thirty μm-thick coronal sections spanning the AP axis of the hippocampus (bregma -1.5-3) were prepared using a Leica cryostat. Free-floating sections were immunolabeled for AT8 using the Mouse-on Mouse Elite peroxidase immunodetection kit (Vector Laboratories, PK-2200). Briefly, endogenous peroxidases were quenched in hydrogen peroxide buffer (PBS + 10% methanol + 3% H_2_O_2_) for 15 mins at 21℃. Tissue sections were washed 2X in PBS, blocked using the mouse IgG blocking reagent for 1.5 hours at 21℃, and washed again 3X with PBS. AT8-biotin (ThermoFisher MN1020B) primary antibody was incubated with the sections at 1:500 dilution in the M.O.M. diluent overnight at 4℃. Sections were washed 5X in PBS, incubated for 30 mins at 21℃ in ABC working solution (Vectastain Elite HRP kit; Vector laboratories, PK-6100), and washed again 3X in PBS. 3,3′-Diaminobenzidine (DAB) working solution (DAB Peroxidase Substrate Kit; Vector Laboratories; SK-4100) was added to the sections until the colour was fully developed (under constant dissection microscope monitoring). Sections were immediately washed 2X in PBS to stop the reaction, mounted onto glass coverslips, and dehydrated successively in 70% ethanol, 95% ethanol, 100% ethanol, and 100% xylene. Coverslips were mounted using Permount (DPX mounting medium) and dried overnight. DAB labelling was assessed every 5^th^ slice (10 slices per animal) at 20X magnification on a Zeiss Axio Scan microscope. AT8 immunolabelling intensity was quantified from ipsilateral (AD EV injected) and contralateral (control EV injected) brain regions (DG, CA2/CA3) using ImageJ.

### Isolation of extracellular vesicles spiked with sarkosyl-insoluble material from AD brain

Sarkosyl-insoluble material enriched with tau PHFs was extracted from 1 g grey matter from the frontal cortex of an individual who had AD based on^13^. Grey matter was homogenised in 9 volumes of ice-cold extraction buffer containing 10 mM Tris-HCl, pH 7.4, 0.8 M NaCl, 10% sucrose, 1 mM EDTA, 0.1% sarkosyl and protease/phosphatase inhibitor cocktail (Pierce) using a FastPrep-24 beat-beating homogeniser (MP Biomedicals). The homogenate was centrifuged at 11,300 revolutions per minute (RPM) for 10 min at 4°C using a Ti45 rotor (Beckman). The supernatant was filtered through a 70 μm pore size cell-strainer and retained on ice. The pellet was resuspended in 4.5 volumes of ice-cold extraction buffer and re-centrifuged at 11,300 RPM for 10 min at 4°C using a Ti45 rotor (Beckman). The supernatant was filtered through a 70 μm pore size cell-strainer and combined with the first supernatant. The combined filtered supernatant was brought to a final concentration of 1% sarkosyl using 25% sarkosyl in water and incubated for 1 h at 21°C with stirring at 100 RPM. Following incubation, the supernatant was centrifuged at 43,800 RPM for 75 min at 4°C using a Ti45 rotor (Beckman). The floating lipid layer and supernatant were discarded and the centrifuge tube was washed twice with 6 mL PBS, followed by one wash with 2 mL PBS, taking care not to disturb the pellet. The pellet was transferred to a Ti70 centrifuge tube (Beckman) in 3 mL PBS and vortexed. The tube was topped up to 23 mL with PBS and centrifuged at 59,300 RPM for 30 min at 4°C using a Ti70 rotor (Beckman). The floating lipid layer and supernatant were discarded and the pellet was transferred to a 2 mL tube in 50 μL PBS per g initial grey matter. The tube was incubated with rocking at 25 RPM for 16 h at 21°C to soften the pellet, before being resuspended by sonication using a Hielscher S26D11X10 Vial-Tweeter- Sonotrode in 0.5 s pulses at 100% amplitude for a total accumulated power of 200 W. The resuspended pellet was transferred to a TLA110 centrifuge tube (Beckman) and centrifuged at 53,000 RPM for 40 min at 4°C using a Ti70 rotor (Beckman). The supernatant was discarded and the pellet was transferred to a 2 mL tube in 50 μL PBS per g initial grey matter and resuspended by sonication using a Hielscher S26D11X10 Vial-Tweeter-Sonotrode in 0.5 s pulses at 100% amplitude for a total accumulated power of 200 W. The resuspended pellet was centrifuged at 10,000 x g for 30 min at 4°C and the supernatant, containing extracted tau PHFs, was retained. EVs from 0.3 g control and AD frontal cortex were isolated as described above, but were floated in a simplified OP gradient consisting of 3 mL step gradients of 40%, 30%, 20%, 10% and 0% OP. Immediately prior to papain digestion, 15 μL of tau PHFs were added to the control EVs to achieve the same amount of tau PHFs that would have been isolated from 0.3 g of AD tissue.

### Tau peptide mapping

Individual tau peptide LFQ intensities were imputed and normalised using the same strategy as the protein dataset, and were presented as log2 values. Peptides were arranged from N to C terminal; amino acids 1-441. Near duplicate peptides, corresponding to missed tryptic cleavages and to post-translationally modified peptides, were summed to represent the total abundance of sequenced peptides at each amino acid site. Peptides were treated as individual peptides if they were more than 50% unique from a neighbouring peptide. A resultant 17 peptides between amino acids 6-406 were used in the analysis.

### Sarkosyl extraction of EVs

EVs suspensions were pelleted by centrifugation at 100,000 x g for 70 min at 4°C using a 70.1Ti rotor (Beckman) and resuspended in 20 volumes (w/v) of extraction buffer containing 10 mM Tris-HCl, pH 7.5, 0.8 M NaCl, 10% sucrose, 1 mM EGTA, and 1% sarkosyl. The resuspended EVs were incubated with rotation for 1 h at 21°C, followed by centrifugation at 100,000 x g for 70 min at 4°C. For single-particle cryo-EM, the sarkosyl-insoluble material was further washed by two rounds of resuspension followed by re-pelleting by centrifugation at 100,000 x g for 70 min at 4°C, first using 25 μL of extraction buffer per gram of initial tissue used, then the same volume of 20 mM Tris-HCl, pH 7.4, 100 mM NaCl. The final sarkosyl-insoluble pellets were resuspended in 25 μL of 20 mM Tris-HCl, pH 7.4, 100 mM NaCl per gram of initial tissue used. For Figure 2E, each EV fraction was resuspended in 30 μL PBS, and pooled into three groups: F1-3, F4-6, and F7-8. Equal volumes of each fraction group were sarkosyl-extracted as above but included a 10,000 x g for 10 mins clearing spin after the sarkosyl incubation. An additional wash in 1% sarkosyl buffer was also performed to clear the pellets of lipids. Final sarkosyl-insoluble pellets were sonicated at 65% amplitude for two minutes in 1X Laemmli sample buffer (BioRad; 1610747) containing 50 mM dithiothreitol (DTT), and heated at 95°C for 5 min prior to electrophoresis.

### Proteinase K protection assay

Directly post-centrifugation, each freshly-prepared EV pellet was resuspend in 30 μL PBS. Fractions F1-3, F4-6, and F7-8 were pooled and brought to 1 ug/µL total protein in PBS. Protease reactions were set up in a total volume of 15 μL with 1.5 μL of the pooled EV sample (1.5 μg total protein), and 1 μL of 15 ng/μL proteinase K (PK) stock (ThermoFisher, EO0491) to maintain a 10 ng PK/1 μg total protein ratio). For EV permeabilisation, 1 μL of 1% Triton X- 100 stock was added. Reactions were incubated at 37°C for 30 min, and stopped by the addition of 15 μL 2X Laemmli sample buffer + 2X HALT protease and phosphatase solution (ThermoFisher, 78440) and heating for 5 min at 95°C. The reactions were resolved by SDS-PAGE and immunoblotted with the tauC antibody at 1:5,000 dilution.

### EV fractionation and carbonate membrane purification

EVs were isolated as described above, however, prior to flotation in the OP gradient, the crude EV pellet was resuspended in 1 mL PBS containing 0.1M DTT and incubated at 21°C for 30 min with end-over-end rotation. DTT-treated EVs were pelleted at 100,000 x g for 70 min at 4°C in a TLA100 rotor (Beckman). The supernatant, containing DTT-labile EV surface proteins, was concentrated to under 100 μL in 0.5 mL 3 kDa molecular weight cut-off centrifugal concentrators (Millipore, UFC500396) (Surface sample in Fig 4C). The pelleted DTT-treated EVs were resuspended in 5 mL of 40% OP; layered beneath a step gradient of 2.5 mL of each 20%, 10%, and 0% OP in Buffer B containing 0.1M DTT; and centrifuged at 200,000 x g for 16 h at 4°C. The top 11 mL and the bottom 2 mL of the gradient were transferred to separate 60 mL 45Ti ultracentrifuge tubes (Beckman), topped up with PBS and centrifuged at 100,000 x g for 70 min at 4°C. The pellet from the 2 mL fraction from the bottom of the gradient was resuspended in 100 μL of TBS (Pellet sample in Fig 4C). The pellet from the 11 mL fraction from the top of the gradient was resuspended in 1.1 mL of hypotonic buffer consisting of 5 mM potassium phosphate; passed through a 26-gauge needle 20 times to lyse EVs and extract luminal proteins; and centrifuged at 150,000 x g for 3 h at 4°C in a TLA100 rotor. The supernatant, containing luminal proteins, was concentrated to under 100 μL in 0.5 mL 3 kDa molecular weight cut-off centrifugal concentrators (Luminal sample in Fig 4C). The pellet, containing EV membranes, was resuspended in 5 mL of 40% OP and layered beneath another step gradient as before, but in the absence of DTT. The gradient was centrifuged and fractionated as before, except that the pellet collected from the 11 mL fraction from the top of the gradient was resuspended in 1.1 mL 0.1 M sodium carbonate buffer, pH 11, and incubated for 30 min at 21°C to extract all but integral membrane proteins from the EV membranes. The carbonate-stripped membranes were collected by centrifugation at 150,000 x g for 3 h at 4°C in a TLA100 rotor. The pellet was subjected to a final OP step gradient without DTT, and the 11 mL fraction from the top of the gradient was pellet at 150,000 x g for 3 hrs, The final pellet was resuspended in 100 μL TBS with protease inhibitors (Membrane sample in Fig 4C). Equal proportions of each compartment subfraction were resolved by SDS-PAGE, and immunoblotted with the tauC antibody at 1:5,000 dilution.

### Cryo-ET

Pooled EV fractions 4–6 isolated from 0.8 g tissue were diluted 1:5 in PBS containing a 1:3 dilution of 10 nm gold-conjugated BSA (BBI solutions), applied to glow-discharged 2/2 μm holey carbon-coated 200-mesh gold grids (Quantifoil) and plunge-frozen in liquid ethane using a Vitrobot Mark IV (Thermo Fisher). Tilt series were acquired using a 300 keV Titan Krios microscope (Thermo Fisher) equipped with a K3 detector (Gatan) and a GIF-quantum energy filter (Gatan) operated with a slit width of 20 eV. Tilt series were acquired at a pixel size of 2.13 Å or 1.89 Å at a target defocus of -3 μm to -6 μm. A dose-symmetric acquisition scheme was implemented in Serial EM over a tilt range of +/-60° at 3° increments^64, 65^. Each tilt received a dose of 3.17 e^-^/Å^2^ fractionated over 8 movie frames, resulting in a total dose of 130 e^-^/Å^2^ per tilt series.

### Tomogram processing and analyses

Gain correction, motion correction, and contrast transfer function (CTF) estimation were performed in Warp^66^. Tilt series alignment using the 10 nm gold-conjugated BSA as fiducials was performed in IMOD^67^. Tomograms were reconstructed in Fourier space at a pixel size of 8 Å or 12 Å; filtered using Wiener-like deconvolution; and denoised using the noise2noise machine learning principle^68^ all in Warp. For filament and EV measurements, IMOD was used to perform gain correction, motion correction, contrast function estimation, and tomogram reconstruction using weighted back-projection at a pixel size of 8.51 Å or 7.46 Å. Filament widths and helical crossover distances were measured manually. 3D representation and segmentation was performed using napari-tomoslice (https://github.com/alisterburt/napari-tomoslice) in Napari^69^.

### Subtomogram averaging

Subtomogram averaging was performed based on^70^, with adaptations for helical reconstruction. Tomograms were reconstructed in Warp at a pixel size of 16 Å to aid in the visualisation of tau filaments. Initial processing steps were performed in Dynamo^71^. Filaments were manually picked using the filament with torsion model and 17,323 subtomograms were extracted using a box size of 64 pixels at an inter-subtomogram distance of 16 Å. An initial reference was created by aligning and averaging a random subset of 250 subtomograms, and was subsequently used to align and average all subtomograms. Subtomograms within 16 Å of one another were reduced to a single subtomogram using the remove_duplicates.m script (https://github.com/teamtomo/teamtomo.github.io/tree/master/walkthroughs/EMPIAR-10164/scripts). The metadata table for the remaining 6,133 subtomograms was converted into the Self-defining Text Archiving and Retrieval (STAR) format using the dynamo2m package (https://github.com/alisterburt/dynamo2m). The filament number (Helical Tube ID) for each subtomogram, specified in column 21 of the Dynamo metadata table, was added to the STAR file using the Starparser package (https://github.com/sami-chaaban/starparser). The subtomograms were then re-extracted in Warp at a pixel size of 6 Å and a box size of 204 pixels. Subsequent processing was performed using helical reconstruction methods in RELION-3.1^72^. An initial model was generated by averaging a random subset of 500 subtomograms using relion_reconstruct. 3D auto-refinement followed by 3D classification were then performed without masking or applying symmetry. This yielded reconstructions corresponding to filament types resembling PHFs and SFs. The helical crossover distances were measured from the reconstructions and used to calculate the helical twists for a helical rise of 4.7 Å. Helical symmetry was then applied in all subsequent steps. 3D classification was repeated to obtain final subsets of subtomograms for PHF and SF types, followed by unmasked and masked 3D auto-refinements. The subtomograms were then re-extracted in Warp at pixel sizes of 3.1 Å (PHFs) and 3 Å (SFs), with respective box sizes of 165 pixels and 170 pixels. 3D auto-refinement was repeated, with C2 symmetry imposed for the PHF type. The final reconstructions were sharpened using the standard post-processing procedures in RELION-3.1, and overall resolutions were estimated at a Fourier shell correlation of 0.143 between the two independently refined half-maps, using phase-randomization to correct for convolution effects of a generous, soft-edged solvent mask. Helical symmetry was imposed using the RELION Helix Toolbox. Rigid-body fitting of models of PHFs from EVs and SFs from primary age-related tauopathy (PDB ID: 7NRS) was performed in ChimeraX^73^.

### Cryo-EM

Sarkosyl-insoluble extracts were centrifuged at 10,000 x g for 10 min at 4 °C. The supernatants were retained and centrifuged at 100,000 x g for 1 h at 4 °C. The pellets were resuspended in 30 μL of 20 mM Tris–HCl, pH 7.4, containing 100 mM NaCl and centrifuged at 3,000 x g for 30 s at 21 °C. The supernatants were retained and applied to glow-discharged 1.2/1.3 μm holey carbon-coated 300-mesh gold grids (Quantifoil) and plunge-frozen in liquid ethane using a Vitrobot Mark IV (Thermo Fisher). Images for the first EV dataset were acquired using a 300 keV Titan Krios microscope (Thermo Fisher) equipped with a K2 detector (Gatan) at the European Synchrotron Radiation Facility (ESRF)^74^. Images for the second EV dataset and the cell dataset were acquired using a 300 keV Titan Krios microscope (Thermo Fisher) equipped with a K3 detector (Gatan) at the MRC Laboratory of Molecular Biology. GIF-quantum energy filters (Gatan) operated at slit widths of 20 eV and aberration-free image shift within the EPU software (Thermo Fisher) were used during image acquisition. Further details are given in Table S2.

### Helical reconstruction

Movie frames were gain-corrected, aligned, dose-weighted and summed using the motion correction program in RELION-4.0^75^. The motion-corrected micrographs were used to estimate the contrast transfer function (CTF) using CTFFIND-4.1^76^. All subsequent image-processing was performed using helical reconstruction methods in RELION-4.0^77^. The filaments were picked manually and reference-free two-dimensional (2D) classification was performed to remove suboptimal segments. Initial three-dimensional (3D) reference models were generated de novo by producing sinograms from 2D class averages as previously described^78^. 3D autorefinements with optimization of the helical twist were performed, followed by two cycles of Bayesian polishing and CTF refinement^75^. 3D classification was used to further remove suboptimal segments, after which the 3D autorefinement, Bayesian polishing and CTF refinement procedures were repeated. The final reconstructions were sharpened using the standard post-processing procedures in RELION-4.0, and overall resolution were estimated at a Fourier shell correlation of 0.143 between the two independently refined half-maps, using phase-randomization to correct for convolution effects of a generous, soft-edged solvent mask^79^. Local resolution estimates were obtained using the same phase-randomization procedure, but with a soft spherical mask that was moved over the entire map. Helical symmetry was imposed using the RELION Helix Toolbox. Further details are given in Table S2.

### Atomic model building and refinement

The published atomic model of PHFs from the frontotemporal cortex of an individual with sporadic AD (PDB 6HRE) was fitted to the final reconstructions using ChimeraX. The fitted models were subsequently refined in real-space using COOT^80^. Rebuilding using molecular dynamics was carried out in ISOLDE^81^. The models were refined in Fourier space using REFMAC5^82^, with appropriate symmetry constraints defined using Servalcat^83^. To confirm the absence of overfitting, the model was shaken, refined in Fourier space against the first half-map using REFMAC5 and compared to the second half-map (Figure S8). Geometry was validated using MolProbity^84^. Molecular graphics and analyses were performed in ChimeraX. Model statistics are given in Table S2.

### Data and material availability

The mass spectrometry proteomics data have been deposited to the ProteomeXchange Consortium via the PRIDE^85^ partner repository with the dataset identifier PXD037708. R Markdown code and data analysis for mass spectrometry proteomics, NTA datasets, and supplemental figures describing the imputation strategy are accessible via: https://github.com/duff-lab-team/AD-EV-characterisation (see the Readme guide). The tomogram shown in Figure 3A has been deposited to the Electron Microscopy Data Bank (EMDB) under the accession number 16064. Single-particle cryo-EM datasets have been deposited to the Electron Microscopy Public Image Archive (EMPIAR) under the following accession numbers: 11300 for the EV dataset and 11301 for the cellular dataset. Single-particle cryo-EM maps have been deposited to the EMDB under the following accession numbers: 16035 for the EV map and 16039 the cellular map. The atomic models have been deposited to the Protein Data Bank (PDB) under the following accession numbers: 8BGS for the EV model and 8BGV for the cellular model. Any other data are available from the corresponding authors upon request.

## SUPPLEMENTAL INFORMATION TITLES AND LEGENDS

**Figure S1:**
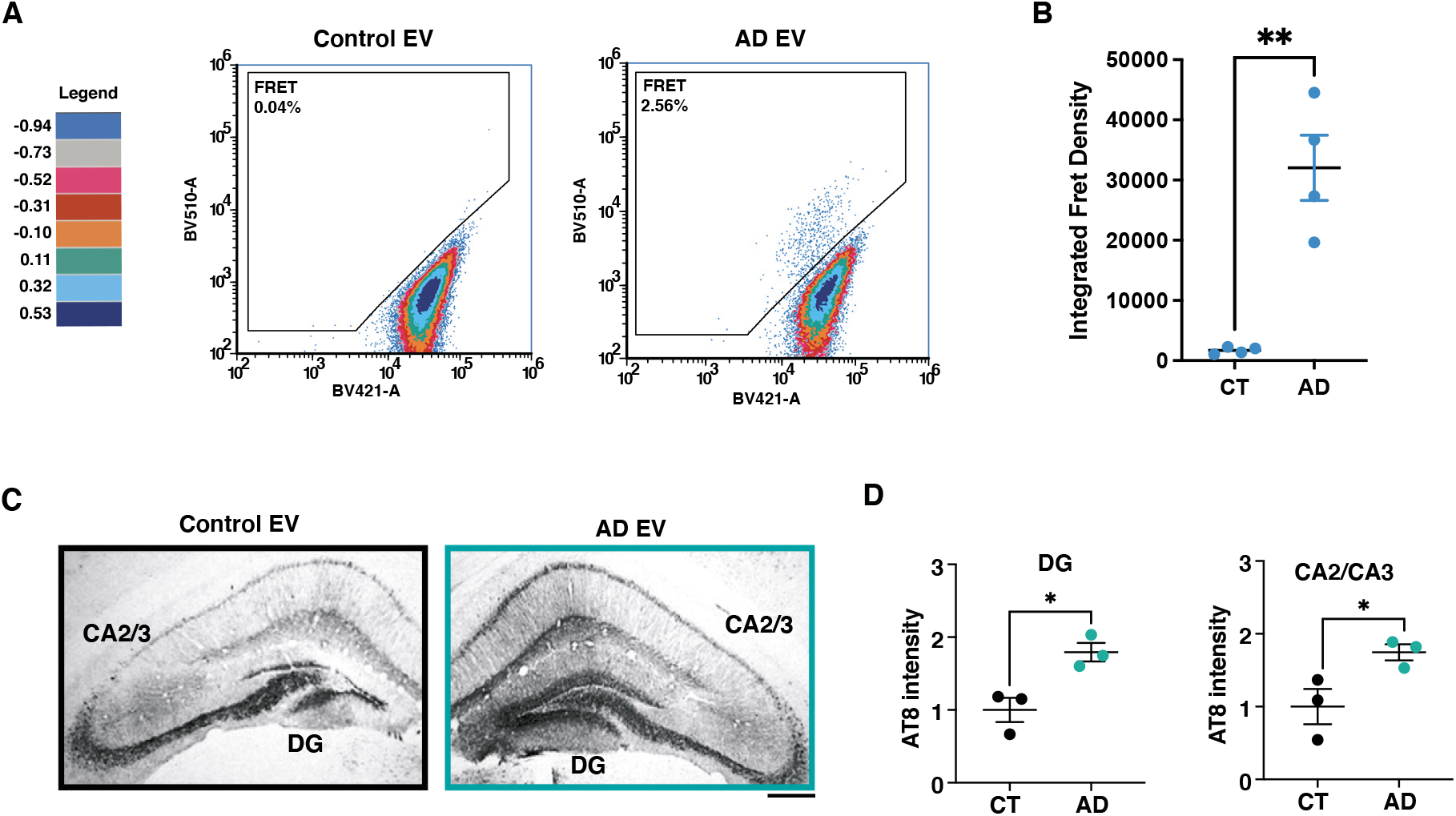
EVs from AD brain induce the accumulation of tau inclusions in the Tau RD P301S FRET Biosensor cell line and in PS19 transgenic mice. (A) Representative FRET gating strategy for the Tau RD P301S FRET Biosensor cell line treated with 3 μg pooled neurologically-normal control and AD EVs. Positive tau seeding is indicated by the presence of BV510 fluorescence signal in the FRET gate. Colour scale indicates log density of measured events. (B) Integrated FRET density (% cells in the FRET gate multiplied by the mean fluorescence intensity of the BV510 signal) of cells incubated for 48 h with 3 μg pooled neurologically-normal control and AD EVs, n=4. Data are means of triplicate technical replicates, two-tailed unpaired t-test. (C) Representative immunohistochemistry images of PS19 mouse hippocampal fields using anti-tau antibody AT8 (phospho-ser202 and -thr205 tau) two-months post stereotaxic injection of pooled EV fractions isolated from neurologically-normal control brain and AD brain into the DG. Scale bar, 100 μm. (D) Quantification of the mean AT8 intensity in the DG and the CA2/CA3 hippocampal fields of PS19 mice injected with EVs from CT and AD brain, n=3, two-tailed unpaired t-test. * = p ≤ 0.05, ** = p ≤ 0.01.

**Figure S2:**
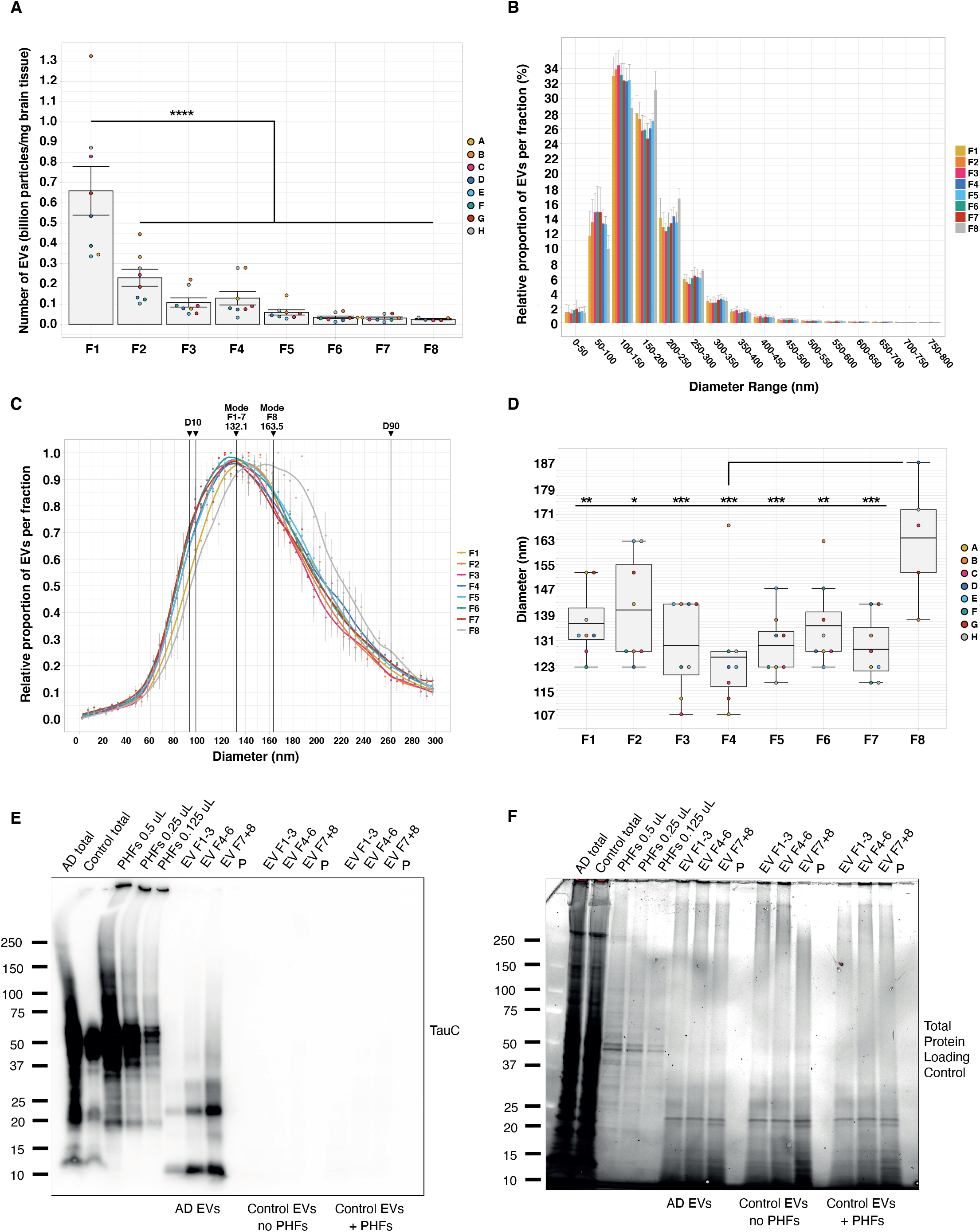
Characterisation and co-purification controls of human brain derived EVs. (A) Nanoparticle tracking analysis (NTA) of total number of EVs per fraction 1–8. Data are expressed as billions of particles per mg brain tissue, one-way ANOVA with Tukey’s post-hoc. (B) NTA analysis of the relative proportion (%) of EVs assigned to the depicted diameter bins for each fraction 1–8 (NTA). (C) NTA analysis of EV diameter for each fraction 1–8 normalised to the mode of each distribution (NTA). D10 (F1-F7: 92.5 nm, F8: 97.5 nm) and D90 (262.5 nm for all fractions) values are means of each value across all fractions. (D) NTA analysis of EV diameters expressed as the mode of each fraction 1–8 for each donor (A–D) n=8 human donors for F1-F7, n=5 for F8, means of triplicate technical replicates were plotted for each donor. Box plot centre lines indicate the mean value, lower and upper hinges represent the first and third quartiles, and whiskers extend from the hinges to a maximum of 1.5X the interquartile range, one-way ANOVA with Dunnett’s post-hoc to F8.* = p ≤ 0.05, ** = p ≤ 0.01, *** = p ≤ 0.001, **** = p ≤ 0.0001. (E and F) Representative immunoblot using antibody TauC (tau repeat region) and corresponding stain-free total protein loading control of AD and control brain homogenates (total); PHFs extracted from AD brain; pooled EV fractions 1–3, 4–6, and 7–8 from AD and control brain, with and without spiked PHFs extracted from AD brain; and the pellet following EV fractionation (P). n=2.

**Figure S3:**
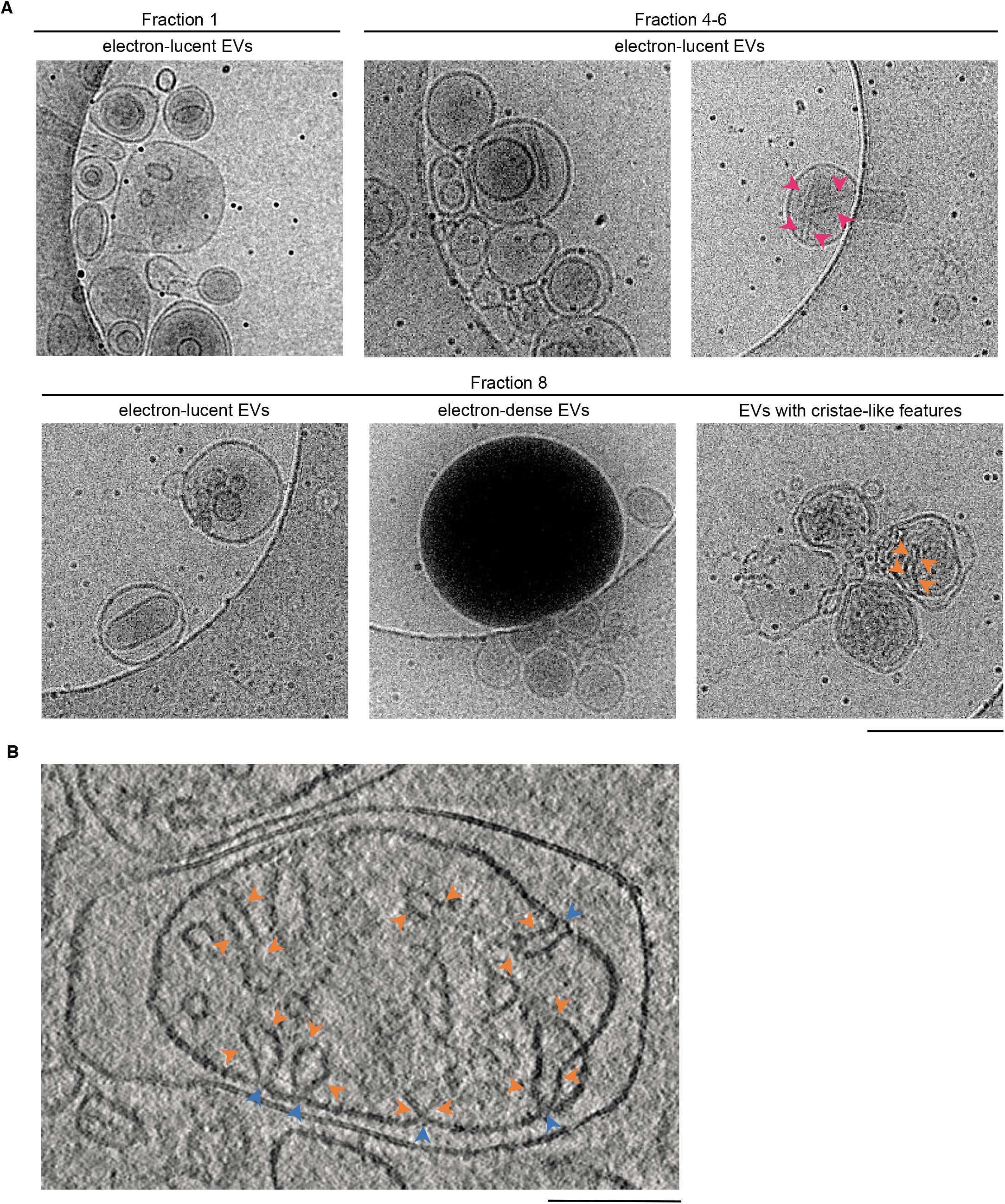
Cryo-EM of extracellular vesicles isolated from Alzheimer’s disease patient brain. (A) Cryo-EM images of morphologically-distinct EV populations in fraction 1, pooled fractions 4–6 and fraction 8. Electron-lucent EVs were observed in all fractions. Electron-dense EVs and electron-lucent EVs with cristae-like features were only observed in fraction 8. EVs associated with tau filaments were only observed in pooled fractions 4–6. Magenta arrows indicate examples of PHF-like filaments and orange arrows indicate examples of cristae-like features. Scalebar, 500 nm. (B) Deconvoluted tomographic slice of an EV from fraction 8 with cristae-like features. Orange arrows indicate cristae-like structures and blue arrows indicate membrane invaginations at the necks of cristae-like structures. Scalebar, 200nm.

**Figure S4:**
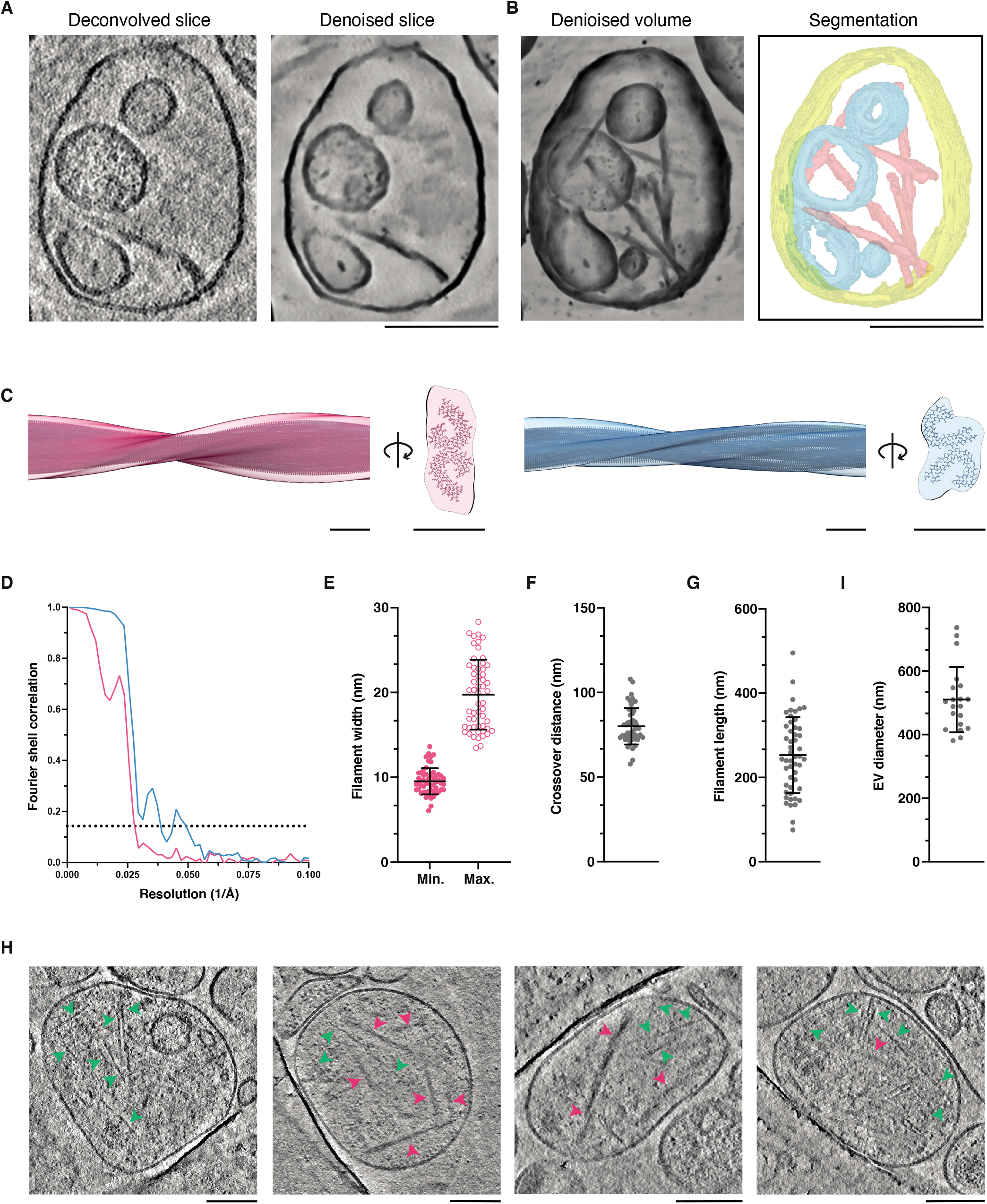
Cryo-ET of extracellular vesicles containing tau PHFs and SFs from Alzheimer’s disease patient brain continued. (A) Deconvolved and denoised tomographic slices of the EV containing tau PHFs and SFs from AD brain shown in Figure 3A. (B) Denoised tomographic volume and segmentation of the EV containing tau PHFs and SFs from AD brain shown in Figure 3A. The limiting membrane is highlighted in yellow, intraluminal vesicles in cyan, and PHFs in magenta. (C) Fit of atomic models (grey) of PHFs (PDB ID 6HRE) and SFs (PDB ID 7nrs) to the subtomogram averaged maps of tau PHFs (magenta) and SFs (blue) in EVs. (D) Fourier shell correlation (FSC) curves for the two independently refined subtomogram averaged half-maps of the PHF-like filament type (magenta line) and the SF-like filament type (blue line). The FSC of 0.143 is shown with a dashed line. (E) Measurements of the minimum and maximum widths of tau filaments in EVs from AD patient brain. n = 55. (F) Measurements of the helical crossover distance (the distance between minimum and maximum widths) of tau filaments in EVs from AD patient brain. n = 50. (G) Measurements of the lengths of tau filaments in EVs from AD patient brain. n = 50. (H) Measurements of the diameters of EVs containing tau filaments from AD patient brain. n, 20. (I) Deconvolved tomographic slices of EVs from AD brain containing tau filaments (magenta arrows) and non-tau filaments (green arrows). Scale bars, 200 nm in (A–C) and 10 nm in (D).

**Figure S5:**
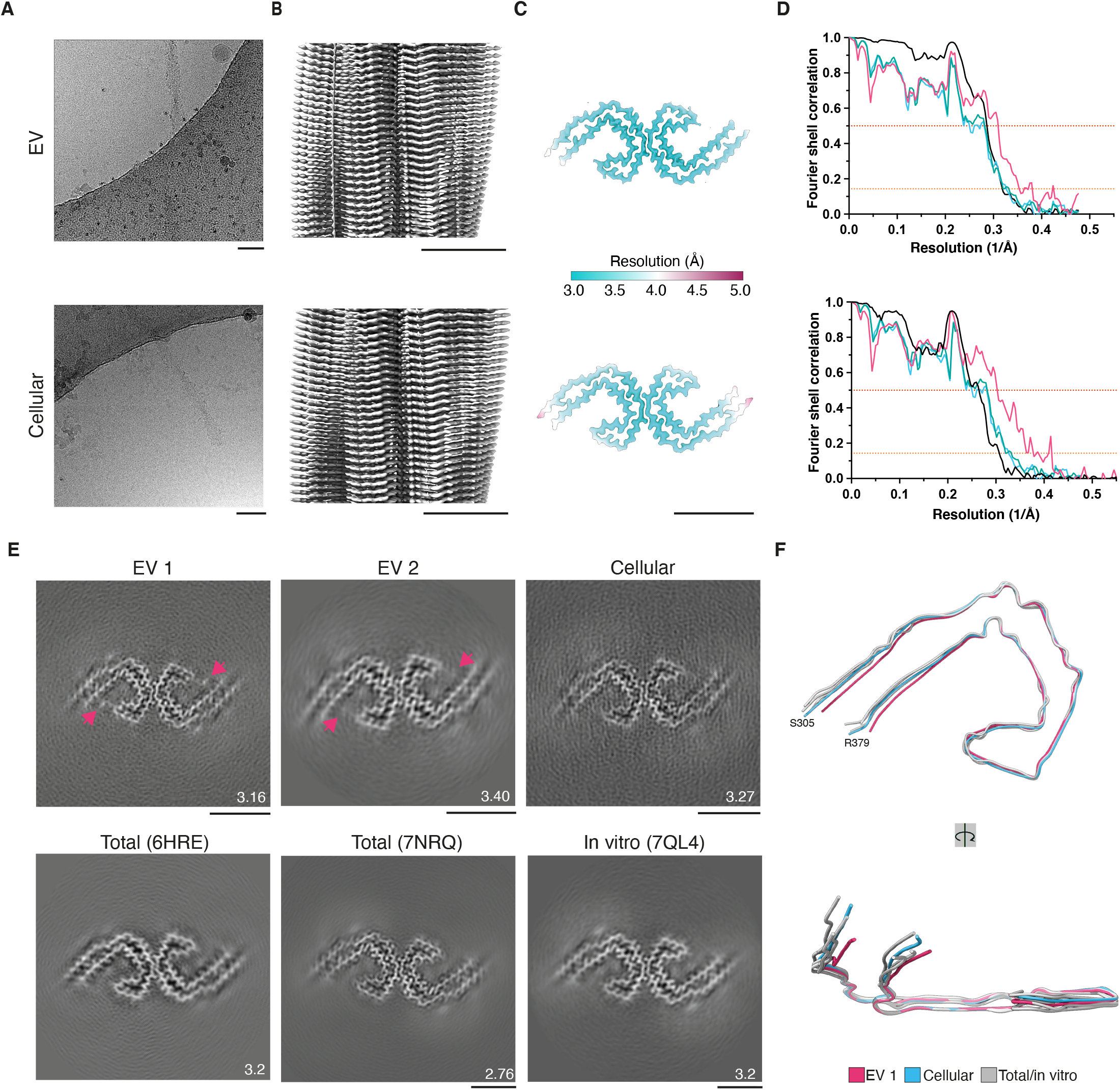
Comparison of cryo-EM structures of PHFs from EV and cellular fractions of Alzheimer’s disease brain. (A) Representative cryo-EM images of PHFs extracted from EVs and from the cellular fraction from AD brain. (B) Cryo-EM maps viewed aligned to the helical axis of PHFs extracted from EVs and from the cellular fraction from AD brain. (C) Local resolution estimates for the cryo-EM maps of PHFs extracted from EVs and from the cellular fraction from AD brain. (D) Fourier shell correlation (FSC) curves for the two independently refined cryo-EM half-maps (black line); for the refined atomic model against the cryo-EM density map (magenta); for the atomic model shaken and refined using the first half-map against the first half-map (teal); and for the same atomic model against the second half-map (cyan) of PHFs extracted from EVs and from the cellular fraction from AD brain. The FSC thresholds of 0.143 and 0.5 are shown with orange and red dashed lines, respectively. (E) Cryo-EM maps of PHFs from EVs from two different individuals, the cellular fraction from one individual, total brain homogenate from two different individuals (PDB IDs 6HRE and 7NRQ), and *in vitro* reconstitution (PDB ID 7QL4), shown as central slices perpendicular to the helical axis. Additional densities in the structures of PHFs from EVs are indicated with arrows. (F) Overlay of the PHF folds from EVs (magenta), the cellular fraction (cyan), and total brain homogenate and *in vitro* reconstitution (PDB IDs 6HRE, 7NRQ and 7QL4) (grey). (A–C and E) Scale bars, 50 Å.

